# Helical transition of the bridge helix of Cas12a is an allosteric regulator of R-loop formation and RuvC activation

**DOI:** 10.1101/2025.01.09.632262

**Authors:** Chhandosee Ganguly, Lindsie Martin, Swarmistha Aribam, Leonard M. Thomas, Rakhi Rajan

## Abstract

CRISPR-Cas12a is widely used for genome editing and biomarker detection since it can create targeted double-stranded DNA breaks and promote non-specific DNA cleavage after identifying specific DNA. To mitigate the off-target DNA cleavage of Cas12a, we previously developed a *Francisella novicida* Cas12a variant (FnoCas12a^KD2P^) by introducing double proline substitutions (K969P/D970P) in a conserved helix called the bridge helix (BH). In this work, we used cryogenic electron microscopy (cryoEM) to understand the molecular mechanisms of BH-mediated activation of Cas12a. We captured five structures of FnoCas12a^KD2P^ at different states of conformational activation. Comparison with wild-type (FnoCas12a^WT^) structures unravels a mechanism where BH acts as a trigger that allosterically activates REC lobe movements by tracking the number of base pairs in the growing RNA-DNA hybrid to undergo a loop-to-helical transition and bending to latch onto the hybrid. The transition of the BH is coupled to the previously reported loop-to-helix transition of the “lid”, essential for opening RuvC endonuclease, through direct interactions of residues of the BH and the lid. We also observe structural details of cooperativity of BH and “helix-1” of RuvC for activation, a previously proposed interaction. Overall, our study enables development of high-fidelity Cas12a and Cas9 variants by BH-modifications.

## INTRODUCTION

CRISPR-Cas systems are adaptive immune systems of bacteria and archaea to protect them from invading genomes^1–6^. While there are different types of CRISPR-Cas systems based on the type of nucleic acid target and protein composition, the fundamental principle is RNA-guided DNA/RNA targeting followed by cleavage of the invader genome. Of the different types, Class 2 systems composed of single multi-domain proteins are widely used for gene editing, including gene therapy^7^. Cas9 (signature protein of the type II system) and Cas12a (signature protein of the type V-A) system are Class 2 proteins widely used for genome applications^8,9^. These proteins are bilobed and target DNA by recognizing a 20 nucleotides (nt) long complementary region between the crRNA-guide and the target DNA. Both Cas9 and Cas12a use protospacer adjacent motif (PAM; a short DNA motif (2-8 nt) flanking the 20-nt region) present in the target to distinguish self *vs.* non-self DNA^10,11^.

Cas9 and Cas12a have similar protein organization features with two lobes, a recognition (REC) lobe and a nuclease (NUC) lobe to perform the functions of RNA-DNA recognition and DNA cleavage, respectively. The two lobes are bridged by a conserved arginine-rich helix, called the bridge helix (BH), and previous studies have shown the importance of BH in RNA-DNA binding and cleavage^12–20^. Each lobe has multiple domains, and in general, both Cas9 and Cas12a undergo conformational changes to accommodate the RNA-DNA hybrid in a groove between the two lobes, even though the exact conformational changes are different^7^. Step-by-step R-loop propagation enables both Cas9 and Cas12a to have conformational check points along the conformational cascade towards achieving the pre-catalytic stage such that formation of the catalytically competent state is tightly related to a full RNA-DNA base pairing in the R-loop (17 nt in Cas9 and 16 nt in Cas12a)^21–23^.

Previous structural^13,22–29^, computational^30,31^, and Förster resonance energy transfer (FRET) studies^17,18,22,32,33^ covering different orthologs have provided an in-depth information on the conformational activation process of Cas12a. The apo-form^27^ of Cas12a is elongated and it transforms into a compact and closed crab-claw shaped structure after binding to crRNA (binary form)^25,29,34^. After binding to the target DNA (ternary form)^13,22–28^, the closed structure opens up to accommodate the growing hybrid in between the two lobes^17,33^. Upon full R-loop formation, there is opening up of the RuvC catalytic pocket through the conversion of a loop form of the “lid” covering the active site to a helix form, which enables passage of single-stranded DNA (ssDNA) into the RuvC active site pocket for cleavage^22,23,26,30^.

Off-target DNA cleavage is a problem in gene editing and different protein engineering strategies are being used to create high-fidelity Cas proteins for genome applications. Previous studies form our lab have shown that the conserved BH of Cas9 and Cas12a can be used for rationale protein engineering to remove off-target DNA cleavage^14,15,20^. Specifically, we targeted a region of the BH that undergoes a loop-to-helical transition in response to RNA/DNA binding [based on comparing different structures (PDBs:5NFV^25^, 5MGA^26^, 5B43^13^, 5ID6^29^, 6I1L^24^, 6I1K^24^, and 5XUS^28^)] and introduced prolines to disrupt the smooth helical transition. The resulting variants showed higher fidelity in DNA cleavage. In this work, we determined the cryogenic electron microscopy (cryoEM) structure of *Francisella novicida* Cas12a (FnoCas12a) with two proline substitutions in the BH (K969P/D970P; referred to as FnoCas12a^KD2P^). The structures unravel a mechanism by which the BH of Cas12a plays an integral part in the smooth transition of the different intermediate states to reach the final active conformation. Specifically, the loop-to-helical transition of BH is related to the number of base pairs in the RNA-DNA hybrid, which allosterically coordinates several downstream processes required to attain a cleavage-competent state, including the opening of the RuvC lid. Since BH is conserved in several types of Cas proteins, as well as in other RNA-binding proteins such as RNA polymerases, our work provides molecular insights on the fidelity of DNA/RNA catalyzing enzymes.

## RESULTS

### FnoCas12a^KD2P^ samples different conformational stages towards the DNA cleavage competent state

To assess how the conserved BH of Cas12a and its helical transitions promote conformational changes needed for DNA cleavage, we determined cryoEM structures of FnoCas12a^KD2P^. We reconstituted FnoCas12a^KD2P^ with its crRNA and a 24 base paired (bp) double-stranded DNA (dsDNA) substrate in the absence of a divalent metal to prevent DNA cleavage. The ternary complex was purified using size exclusion chromatography (SEC) and the peak corresponding to the ternary complex, peak 2 (Supplementary Fig. 1) was used to make cryo-grids. We got consistently better distribution of the particles when we used graphene oxide coated grids^35,36^. An *ab initio* 3D map at ∼2.96 Å was created from ∼400K particles and the map possessed the typical crab-claw shape as the wild-type FnoCas12a (FnoCas12a^WT^, PDB: 6GTG^22^). 3D variability analysis created 6 subclasses with an almost equal distribution of particles (14% to 18%, Supplementary Fig. 2). Five of the subclasses ranged in resolution from 3.2 Å to 4 Å (Supplementary Fig. 3, Supplementary Table 1) and these were used for model building and analysis. These five structures have differences in domain placements and represent conformational transitions needed to activate Cas12a for DNA cleavage (Fig. 1).

**Figure 1:**
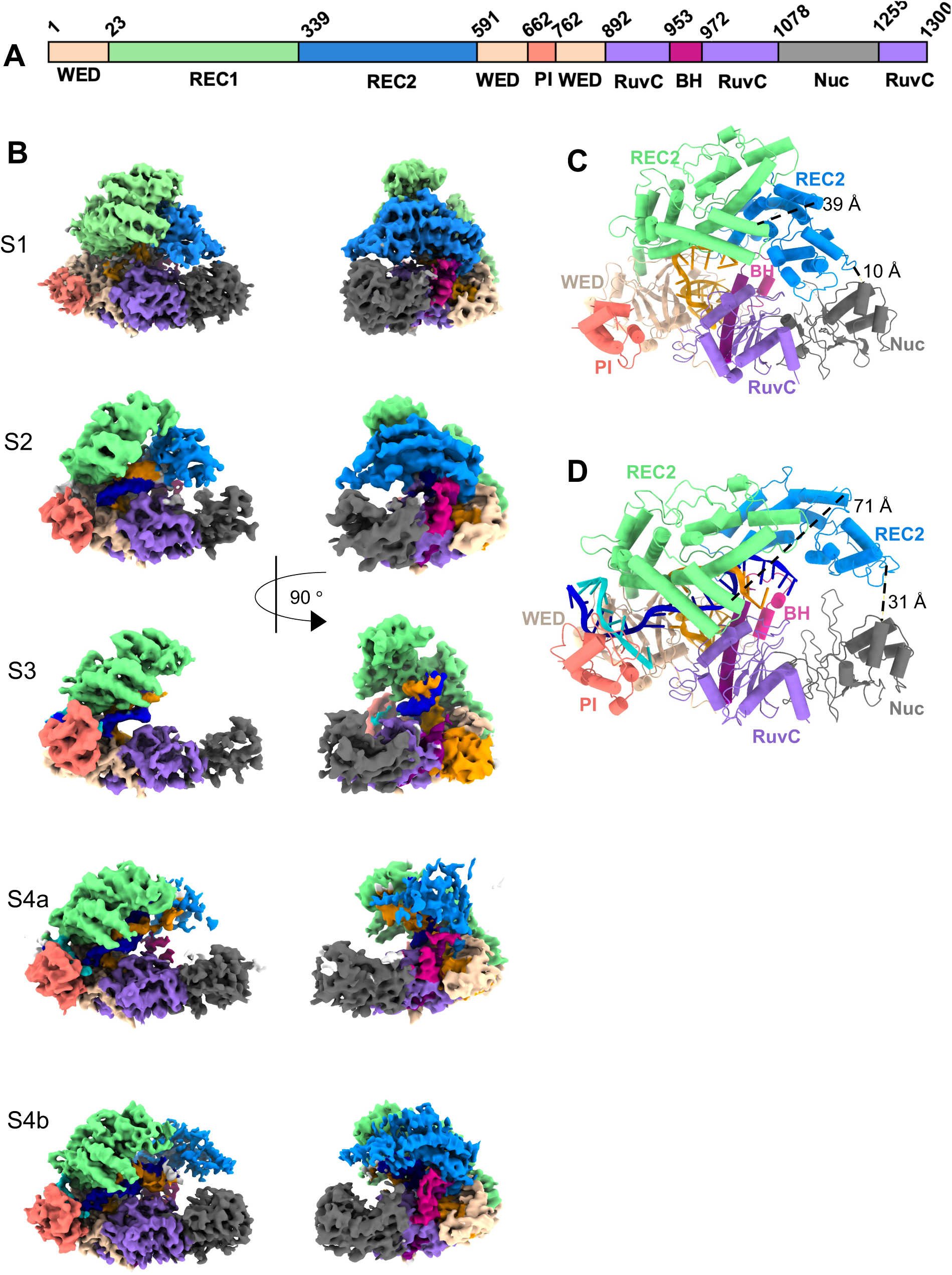
Conformational states of Cas12a towards attaining the pre-catalytic state. **(A)** Schematic representation of the domain architecture of Cas12a. The domain colors are same as this scheme throughout the manuscript. **(B)** CryoEM maps of the different conformational intermediates of FnoCas12a^KD2P^ (S1 to S4b) displayed in two different orientations. The major change is in the positioning of REC1 and REC2 domains between the different states. Model built to fit the map of state 1 **(C)** and state 4b **(D)**. Between the states S1 and S4b, the complex transitions from a closed to an open conformation, going from a distance of 10 Å between the REC2 and Nuc domains in S1 to 31Å in S4b. The distance between the REC1 and REC2 domains also changes from 39 Å to 71 Å, respectively in states S1 and S4b (also see **Supplementary Fig. 3**).

The maps and models of FnoCas12a^KD2P^ ternary complex from this study were compared with those of FnoCas12a^WT^ from a previous study (Supplementary Table 2) that used single-molecule FRET (smFRET) analysis to identify the different conformational states (intermediate states I1-I5) towards attaining the pre-catalytic conformation^22^ as well as that of an *Acidaminococcus sp.* (As) Cas12a structure series that captured different conformations at different lengths of the R-loop formation^23^. We named the different structures of FnoCas12a^KD2P^ complex as S1 to S4b (Fig. 1B). These structures provide direct evidence on the role of BH in orchestrating the conformational cascade to activate Cas12a for cleavage, including the opening of the lid covering the RuvC active site through direct interactions with the BH residues.

Comparison of the map features of all the five states revealed that the major difference between them is in the movements and visibility of the REC1 and REC2 domains, which was previously shown to be important to open the closed binary complex of FnoCas12a^WT^ to accommodate the RNA-DNA hybrid in between^22,23^. Notably, in state S1, the REC2 domain of the REC lobe is placed over the Nuc domain giving it a closed conformation (Fig. 1C, Supplementary Table 3) and the PAM-interacting (PI) domain remains open compared to the other states, due to some disordered residues. S1 did not show continuous stretch of density for DNA, but there is residual density for positions 4 to 6 of the TS DNA (Supplementary Fig. 4). An analysis of the seed region of the crRNA shows that there is preordering of this region similar to the binary complex (PDB: 5NG6^25^, Supplementary Fig. 4), indicating that the BH substitutions do not impact binary complex formation. These factors suggest that the S1 state represents initiation of the ternary complex formation where the complex is in very early stages of PAM recognition.

In state S2, the ternary complex transitions toward R-loop formation. The protein undergoes a closed to open transition, where the REC1 and REC2 domains begin to move away from and upward relative to the NUC lobe to accommodate the R-loop (Supplementary Tables 3 and 4). During the transition of S1 to S2, the REC1 domain undergoes ∼14° rotation with respect to a horizontal axis passing through the PI and Nuc domains (referred to as NUC-lobe axis here-after, Supplementary Fig. 5) and a translation of ∼14 Å (distance between PI and REC1 domains). In contrast, the REC2 undergoes only a minor rotation of ∼1° away from the NUC-lobe axis (Supplementary Fig. 6A, Supplementary Tables 5A and 5B). We observe 8 nt bp between crRNA and TS DNA (Supplementary Table 6). However, due to the disordered PI domain, no density for the PAM sequence is visible. The single stranded non-target strand (NTS) is also not visible. S2 of FnoCas12a^KD2P^ resembles the state I2 of FnoCas12a^WT^ (PDB: 6GTD^22^; RMSD 3.6 Å, Supplementary Table 2).

In state S3 we observe a complete absence of density for the REC2 domain due to increased flexibility, and hence this domain was omitted from the S3 model (Supplementary Fig. 6B). The movements of REC1 and REC2 domains in S3 increases the accessibility for the RNA-DNA hybrid. In S3, 8 nt of the TS are base paired with the crRNA-guide region (Supplementary Table 6). It is interesting to note that the PI domain is ordered in S3, after 8 nt of base pairing has occurred, and not in S2. This may correlate with the target DNA search mechanism of Cas12a, where establishing a critical amount of base pairing between the TS and crRNA-guide may trigger PI ordering as a cue to further proceed with R-loop formation rather than dissociation and search for another potential DNA target^37^. The S3 of FnoCas12a^KD2P^ appears to be an intermediate state between I2 (PDB: 6GTD^22^) and I3 (PDB: 6GTE^22^) of FnoCas12a^WT^. While REC2 domain was flexible in the FnoCas12a^WT^ structures, none of them was captured with the complete absence of density for the REC2 domain. Similar states with a dislodged REC2 without density was captured for AsCas12a with 8 (PDB: 8SFI^23^) and 10 (PDB: 8SFJ^23^) bp RNA-TS DNA hybrid during R-loop propagation, pointing to large movements needed for REC2 to accommodate the growing R-loop.

The remaining two states of the FnoCas12a^KD2P^ structures are very close in conformations, without significant differences to place them in an order to reach the active state. These states closely resemble the intermediate state I3^22^ of FnoCas12a^WT^ (S4a: 5.2 Å RMSD and S4b: 4.3 Å RMSD, Supplementary Table 2) and the two intermediate states of AsCas12a with 15 and 16 bps of R-loop formation (PDB: 8SFL and 8SFN^23^, respectively, Supplementary Table 2). In S4a, we see less well-defined density for REC2 (similar to 15 bp complex of AsCas12a), with density pattern showing stretched features consistent with the high flexibility of this region. With the current positioning of REC2 in S4a, we observe a rotation of ∼17° upwards of the NUC-lobe axis and a translation ∼14 Å away from the Nuc domain, compared to that in S2 (Supplementary Table 5B) and the distance between REC1 and REC2 increases by ∼77 Å (Supplementary Table 4). S4a has the most visible length of RNA-DNA hybrid of all the five states with 11 bp hybrid between crRNA and TS DNA and 8 nt for the NTS. State S4b has the highest resolution (3.21 Å) among all the five states, with clear density for the REC2 domain (Fig. 1). The REC2 domain rotates downwards to the central cavity compared to its position in S4a (Supplementary Table 3, Supplementary Fig. 6C). In S4b, we see 9 bp crRNA: TS DNA hybrid and 9 nt for the NTS (Supplementary Table 6).

Compared to S1, we observe that the REC1 domain moves a maximum of 14° away from the NUC-lobe axis in S2 and is stable for the rest of the states (Supplementary Table 5A, Supplementary Fig. 7). Comparatively, REC2 moves to a maximum of 18° in S4a compared to S1 and then moves slightly closer to the NUC-lobe axis in S4b (Supplementary Table 5B, Supplementary Fig. 7). As the REC lobe opens, the R-loop grows in between the two lobes.

### Sampling of the pre-catalytic state is reduced in FnoCas12a^KD2P^

We compared the conformational states of FnoCas12a^KD2P^ with that of FnoCas12a^WT^ from different studies (Supplementary Table 2). State S1 appears to be a conformation in between the binary (PDB: 5NG6^25^) and the state I1 (PDB: 6GTC)^22^, which is the initial stage of ternary complex formation based on smFRET studies. The state S2 of FnoCas12a^KD2P^ is similar to state I2 (PDB: 6GTD^22^) due to the similar positions of REC1 and REC2 domains (Supplementary Table 4). There is a ∼7° rotation of REC2 towards the NUC-lobe axis in I4 compared to that in I3 (Supplementary Table 5B). The new states of S3 and S4a of FnoCas12a^KD2P^ likely represent intermediate states between I2 and I3^22^ that were captured during this rotation of REC2, supported by the weak density for REC2 in these states. While in state S4b there is ordering of the REC2 density, it has only 9-nt base pairing with the crRNA guide visible, compared to 11 nt base pairing in S4a (Supplementary Table 6). Overall, states 4a and 4b appear to be very transient states towards attaining a pre-catalytic state (PDB: 6GTG^22^). While we captured all the different states with an equal probability, we did not observe a pre-catalytic state where the RNA-DNA hybrid is ordered with more visibility for NTS closer to the RuvC active site pocket, and also with the critical loop-to-helical transitioning of the “lid” region to open the catalytic pocket. An impaired BH due to the double proline substitutions appears to reduce the efficiency of conformational transitions to reach the catalytically competent state, which also enabled capturing of sub-intermediate states that were not previously observed. This observation supports the lower rate constant that was observed for DNA cleavage by FnoCas12a^KD2P^ compared to that by FnoCas12a^WT^ (3-fold reduction in on-target DNA cleavage^14^).

### Cooperativity in the conformational transitions of BH and helix-1 is critical for Cas12a’s activity

It was previously reported that BH and helix-1 of RuvC domain undergoes coordinated conformational transitions while transforming from the binary to the ternary complex. A comparison of the R-loop initialization (PDB:6GTC, RMSD between 6GTC^22^ and PDB: 5NG6^25^ is 1.15 Å) and 20 bp R-loop (PDB: 6GTG^22^) structures show that there is melting of helix-1 upon partial R-loop formation, followed by elongation of the BH by 3 turns. In addition, the BH-end (D970) bends by 28° and moves ∼20 Å towards the TS-RNA hybrid after full R-loop formation compared to the position in the R-loop initialization structure (Figs. 2A-2C, Supplementary Fig. 8, and Supplementary Tables 7-8). Comparing the structure of Cas12a’s BH in the different states of FnoCas12a^KD2P^ in our study with that of wild-type, we find that the loop-to-helical transition of the BH is impaired due to the proline substitutions. The impairment in the formation of an elongated BH keeps it positioned at a distance of ∼20 Å away from the TS-RNA hybrid in both S4a and S4b (compared to ∼17 Å in 6GTC^22^ and ∼6 Å in 6GTG^22^, Supplementary Table 8). Another important conformational change towards activation of Cas12a is the insertion of the conserved W971 into the REC2 hydrophobic pocket (composed of amino acids Y579, K527, K524, R583 of FnoCas12a^WT^)^13,17,26^ to enable REC2 movements. In FnoCas12a^KD2P^ structures, we see insertion of W971 into the REC2 hydrophobic pocket in S1 and S4b (S2 and S4a does not show high resolution in this area to determine side chain position). However, the residues W971 and Y579, which shows an interaction in FnoCas12a^WT^ (∼4 Å) remains far in S1 and S4b (∼10 Å) (Figs. 2D-2G). It can also be envisioned that W971 as part of a completely elongated BH will be sturdier than being present in an impaired BH. These observations indicate that impairing the loop-helical transition of BH potentially prevents sampling of the activated state in our cryoEM classes due to absence of an elongated and bend BH to latch on to the RNA-DNA hybrid and the potential hindrance in coordinating the REC2 movement by W971.

**Figure 2.**
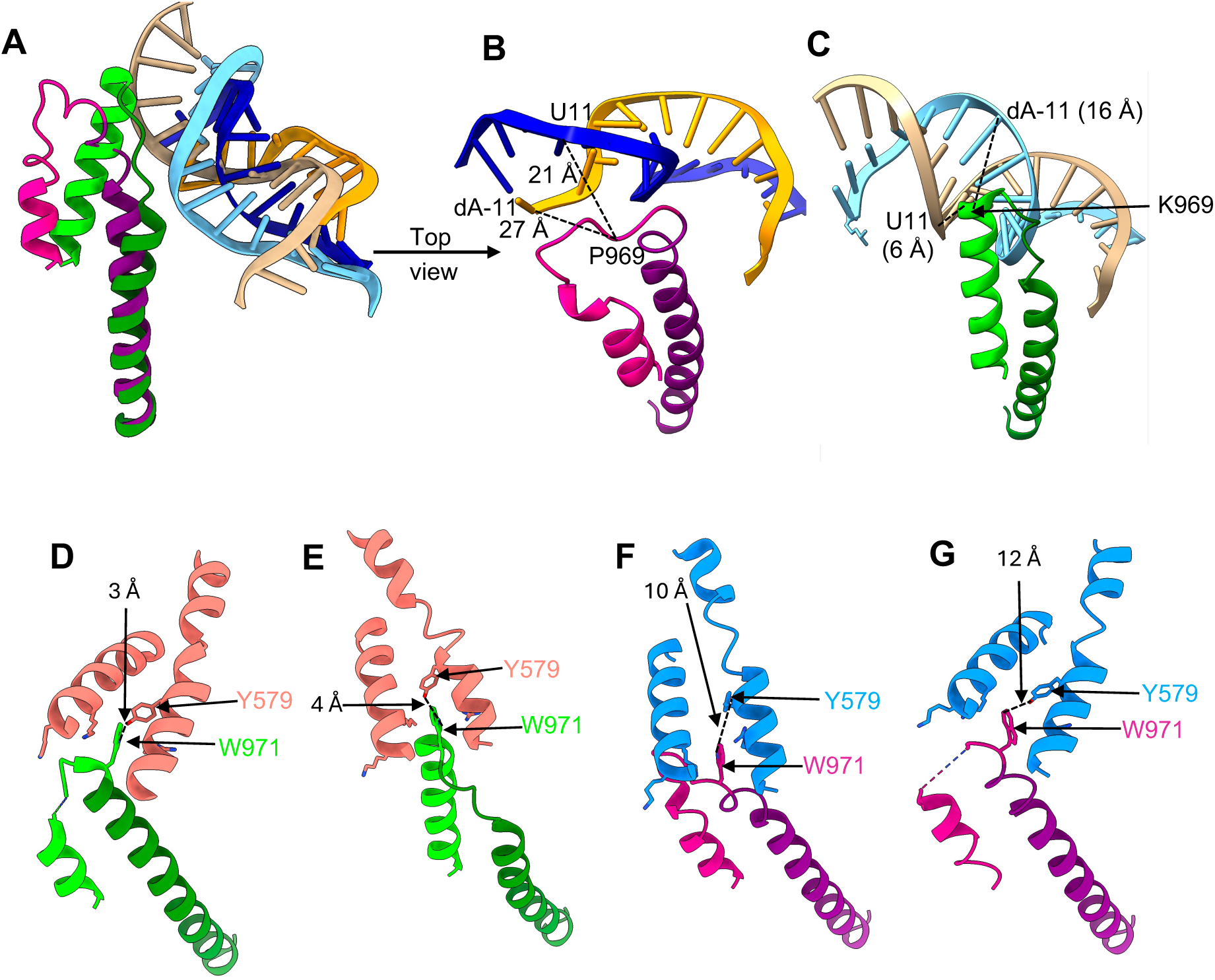
BH transitions are important for R-loop propagation and REC domain movements. **(A-C)** Impaired BH disrupts interactions with the RNA-DNA hybrid. **(A)** An overlay of the FnoCas12a^WT^ structure (green) with 20 nt-long base pairing between the RNA-DNA hybrid (sand and cyan, PDB: 6GTG^21^) with state S4b of FnoCas12a^KD2P^ (BH: magenta, helix-1: purple, RNA-DNA hybrid: orange-blue). BH undergoes loop-to-helical transition and bending to latch on to the RNA:DNA hybrid in FnoCas12a^WT^. A similar transition is absent in FnoCas12a^KD2P^. The RNA-DNA hybrid structure is distorted and far from the BH in FnoCas12a^KD2P^ **(B)**, while the hybrid has an expected helical structure and is closer to the BH in FnoCas12a^WT^ **(C)**. A twist in the RNA-DNA backbone path occurs between the two structures at position 11 of the RNA-DNA hybrid due to the interaction of the BH region undergoing the loop-to-helical transition. **(D-G)** Impact of BH in W971 insertion. The conserved W971, immediately following the proline substitutions, inserts into the REC2 hydrophobic pocket and interacts closely (∼4 Å) with Y579 in both the binary **(D**, PDB: 5NG6) and full R-loop states **(E**, PDB: 6GTG^21^) in FnoCas12a^WT^. The impaired BH of FnoCas12a^KD2P^ impacts this interaction in S1 **(F)** and S4b **(G)** where W971 and Y579 are positioned at ∼10 Å away.

### Impaired BH restricts major conformational checkpoints during R-loop formation

A series of conformational changes including the lysine helix-loop (LKL) of the PI domain, the helix-loop-helix (HLH) segment, the finger region, and the REC-linker of the REC1 domain were shown to be essential for R-loop propagation (Fig. 3). As observed in FnoCas12a^WT^ complexes, in S3 (Fig. 3A), S4a, and S4b, the LKL helix inserts into the double stranded PAM sequence and initiates PAM unzipping (PDB:5MGA^26^). However, the interactions are not observed in S1 and S2 due to disordered PI domain. In the next step, the HLH segment undergoes a positional shift to accommodate the growing R-loop (Supplementary Fig 9, Supplementary Table 9). In our analysis of FnoCas12a^KD2P^ structures, we observed a gradual movement of the HLH segment from S1 to S4b (∼14 Å) (Supplementary Table 9). This movement is the same as that observed in I1 (PDB:6GTC^22^) to I3 (PDB:6GTE^22^) and the position of HLH segment in S4b remains farther away from I4 (PDB: 6GTG^22^) by ∼8 Å (Supplementary Table 9, Fig. 3B and Supplementary Fig. 9).

**Figure 3:**
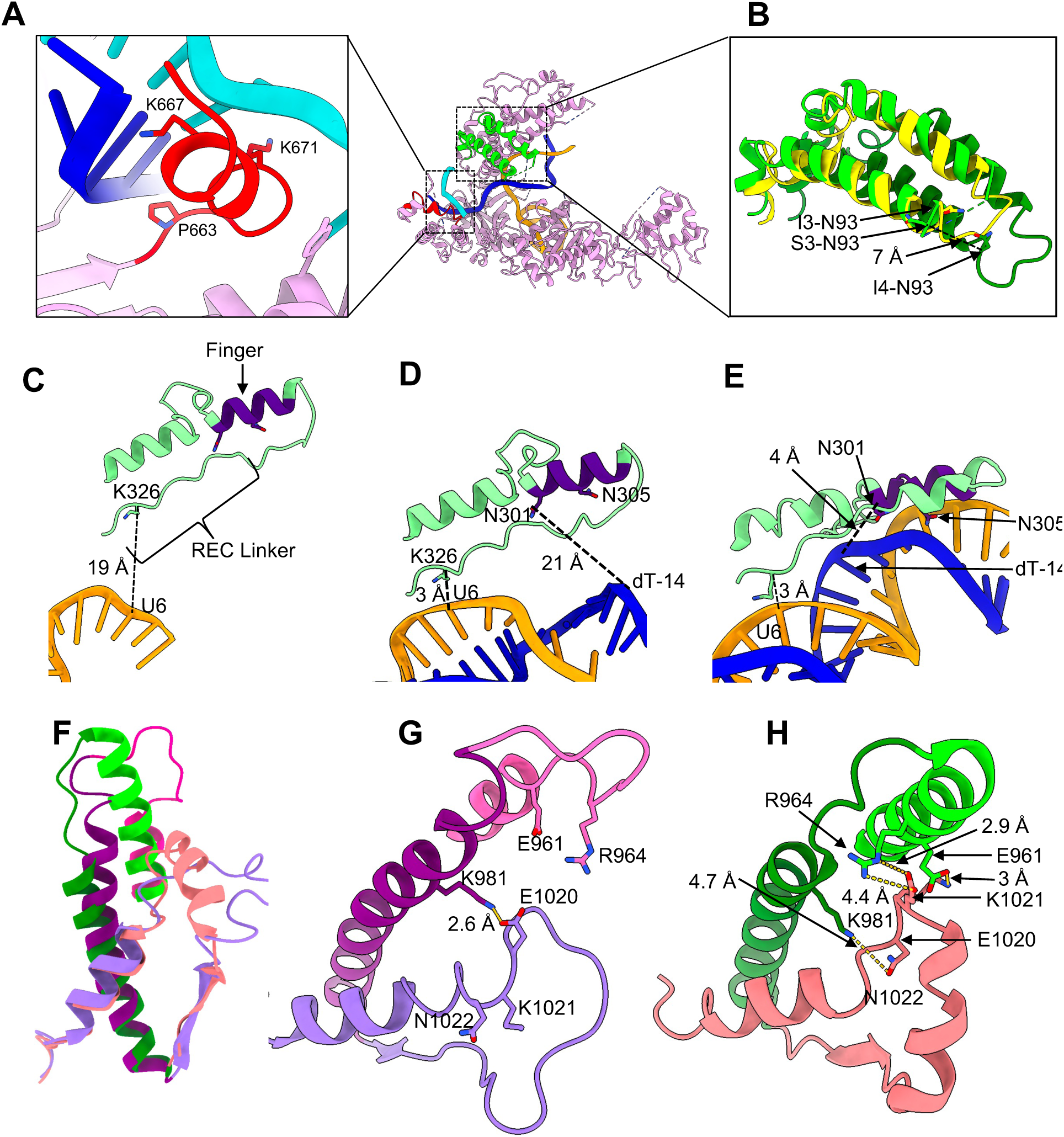
Comparison of the different elements needed for activation of Cas12a in FnoCas12a^KD2P^ and FnoCas12^WT^. Positioning of LKL (red) of the PI domain **(A)** and HLH (lime green) of the REC1 domain **(B)** in S3 of FnoCas12a^KD2P^. The LKL region shows similar interactions through K667, K671 and P663 as in FnoCas12a^WT^ to promote PAM unzipping. **(B)** The HLH domain of S3 is placed in a similar position as the corresponding I3 state of FnoCas12a^WT^, while it is 7 Å farther away compared to its position in the pre-catalytic I4 state of FnoCas12a^WT24^. **(C-E)** Comparison of the REC-linker (green) and finger region (purple) movements in state S1 **(C)**, S4b **(D),** and in FnoCas12a^WT^ **(E** PDB: 6GTG^21^). While the REC-linker movements are similar in both proteins, finger region of FnoCas12a^KD2P^ remains farther at the PAM distal end of the R-loop compared to its position in FnoCas12a^WT^ pre-catalytic state (also see **Supplementary Fig 10**). **(F-H)** Impact of BH in RuvC lid conformations. **(F)** An impaired BH delays the loop-to-helical transition of the RuvC lid that is needed to open the active site pocket for DNA access. (**G)** State S4b of FnoCas12a^KD2P^ maintains pre-catalytic stage interactions of the lid with the helix-1. (**H)** In comparison, in the pre-catalytic stage of FnoCas12a^WT^ (PDB: 6GTG^21^), new interactions are formed between residues of the BH that undergo loop-to-helical transition and the lid. K981 of the helix-1 forms a new interaction with the RuvC (N1022), acting like an anchoring base for the conformational transitions. Colors for panels **E** to **G**: S4b: BH in deep pink, helix-1 in purple, and RuvC lid in medium purple 6GTG: BH in lime green, helix-1 in forest green, and RuvC lid in salmon.

The REC-linker and finger regions were shown to sense the crRNA:TS hybrid formation through PAM-proximal and PAM-distal interactions, respectively^22^. The movement of the REC-linker across the states in FnoCas12a^KD2P^ remains similar to FnoCas12a^WT^. The REC-linker remains positioned away from the crRNA in S1, similar to the R-loop initialization complex (PDB: 6GTC)^22^ and subsequently shifts closer to the crRNA (G5 to C7, PAM-proximal region, similar to PDB: 6GTG^22^), with R-loop propagation from S2 to S4b (Figs. 3C-3E, Supplementary 10 and Supplementary Table 10). The finger region moves closer to the PAM-distal region of the crRNA:TS DNA hybrid during the transition from states S1 to S4b. Despite this movement, docking of the finger region onto the crRNA at positions 15–17 nt, which was identified as a delayed checkpoint towards the catalytic activation of Cas12a^22,23^ is not observed in the structural transitions that were captured for FnoCas12a^KD2P^. Overall, while the interaction of the REC-linker at the PAM-proximal side is intact in FnoCas12a^KD2P^, the interaction of finger at the PAM-distal side is impaired.

### BH is an allosteric regulator that connects RNA-DNA hybrid formation to opening of the lid of the RuvC active site

Previous studies have shown that the lid (residues 1008 to 1021 in FnoCas12a) that covers the active site pocket of the RuvC undergoes a loop-to-helical transition to expose the pocket for DNA access. A comparison of the different states of FnoCas12a^KD2P^ with that of FnoCas12a^WT^ provides interesting insights on the cooperative action of BH and the lid to open the RuvC active site for DNA access.

Regions E961-D970 of the BH in the binary FnoCas12a structure (PDB: 5NG6^25^) undergoes loop to helical transition in response to full R-loop formation (PDB: 6GTG^22^). Similar to AsCas12a^23^, in FnoCas12a the residue E1020 of the lid region interacts with K981 of helix-1 until the 15 bp R-loop formation and stabilizes the loop form of the lid^22,30^. However, analysis of FnoCas12a^WT^ 20 bp R-loop structure (6GTG^22^) shows disruption of E1020-K981 interaction and formation of new interactions between E1020, K1021 and N1022 of the lid with R964 and E961 of the BH and K981 of helix-1, respectively, which enables ordering of the lid-helix (Figs. 3F-3H). Thus, it appears that the helical transition of the BH in response to DNA binding translates to the helical transition of the lid region, which in turn opens the active site pocket of the RuvC for DNA access. We propose the BH structural changes, which is linked to the R-loop length, to be the initiating trigger to open the RuvC active site pocket.

An analysis of the BH and lid conformations in our FnoCas12a^KD2P^ structures provide support for this proposed mechanism. In all the five states that we observed, the lid is in a loop form and blocks the RuvC active site pocket (Fig. 11). In S1 and S4b, we see full density for the lid-loop region, while in S2, S3, and S4a, the density is absent for residues 1009 to 1017 (Supplementary Fig. 11A). In all the five states that we observed, the BH is still in the loop form and has not transitioned into the helical form in response to DNA binding. The introduction of double proline substitutions in the BH-loop hinders its helical transition, which in turn obstructs the helical formation of the lid (Supplementary Figs. 8 and 11). This is supported by our previous experimental work that showed a 3-fold reduction in the cleavage of matched (on-target) plasmid DNA by FnoCas12a^KD2P^ compared to that of FnoCas12a^WT14^.

### BH conformational change is essential for R-loop formation and stability

A comparison of the RNA-DNA hybrid in the different states of FnoCas12a^WT^ and FnoCas12a^KD2P^ shows that the BH is essential for RNA-DNA hybrid propagation and proper base pairing. An analysis of the distance evolution between the BH and the RNA/DNA is shown in Supplementary Table 8. K969 is at a distance of 34 Å from U4 of the crRNA in the binary structure of FnoCas12a^WT^ (PDB: 5NG6^25^) and the corresponding distance is 34 Å between P969 with A4, showing similar positioning of the crRNA in S1 of FnoCas12a^KD2P^. In the pre-catalytic state of FnoCas12a^WT^, the distance between the RNA/DNA hybrid gets closer (∼6 Å with U11) when the R-loop has a full 20-nt RNA-DNA base pairing^22^ (Figs. 2 and Supplementary Fig. 8, and Supplementary Table 8). In S4a and S4b of FnoCas12a^KD2P^, the RNA-DNA hybrid remains farther (20 Å and 27 Å respectively) from the BH due to the inability of the BH to undergo the loop-to-helical transition, which prevents its extension towards the hybrid (Figs. 2, Supplementary Fig. 8, and Supplementary Table 8).

In FnoCas12a^KD2P^ states, RNA:TS DNA base pairing pattern is impacted by the BH. In S2, a region of the PI domain (S695-I714) is disordered, and there is no clear density for the PAM sequence of the DNA, compared to clear visibility of 4 nt PAM and 8 nt RNA:DNA hybrid in 6GTC. In S2 and S3, after dG(-7), the R-loop exhibits unstable base pairing. However, for S4a and S4b of FnoCas12a^KD2P^, base pairing is stable until position dA(-11) and dA(-10) of TS DNA with the corresponding crRNA-guide complementary position, with the remaining visible nt (4 nt in S4a and 3 nt in S4b) in a single stranded condition (Fig. 4) . We observe stacking and distortion of the single-stranded form of the crRNA and the TS DNA in FnoCas12a^KD2P^ structures, beyond the base paired region. This indicates that the BH conformational change is essential for progressive R-loop formation and to stabilize the base pairing of the RNA-DNA hybrid.

**Figure 4:**
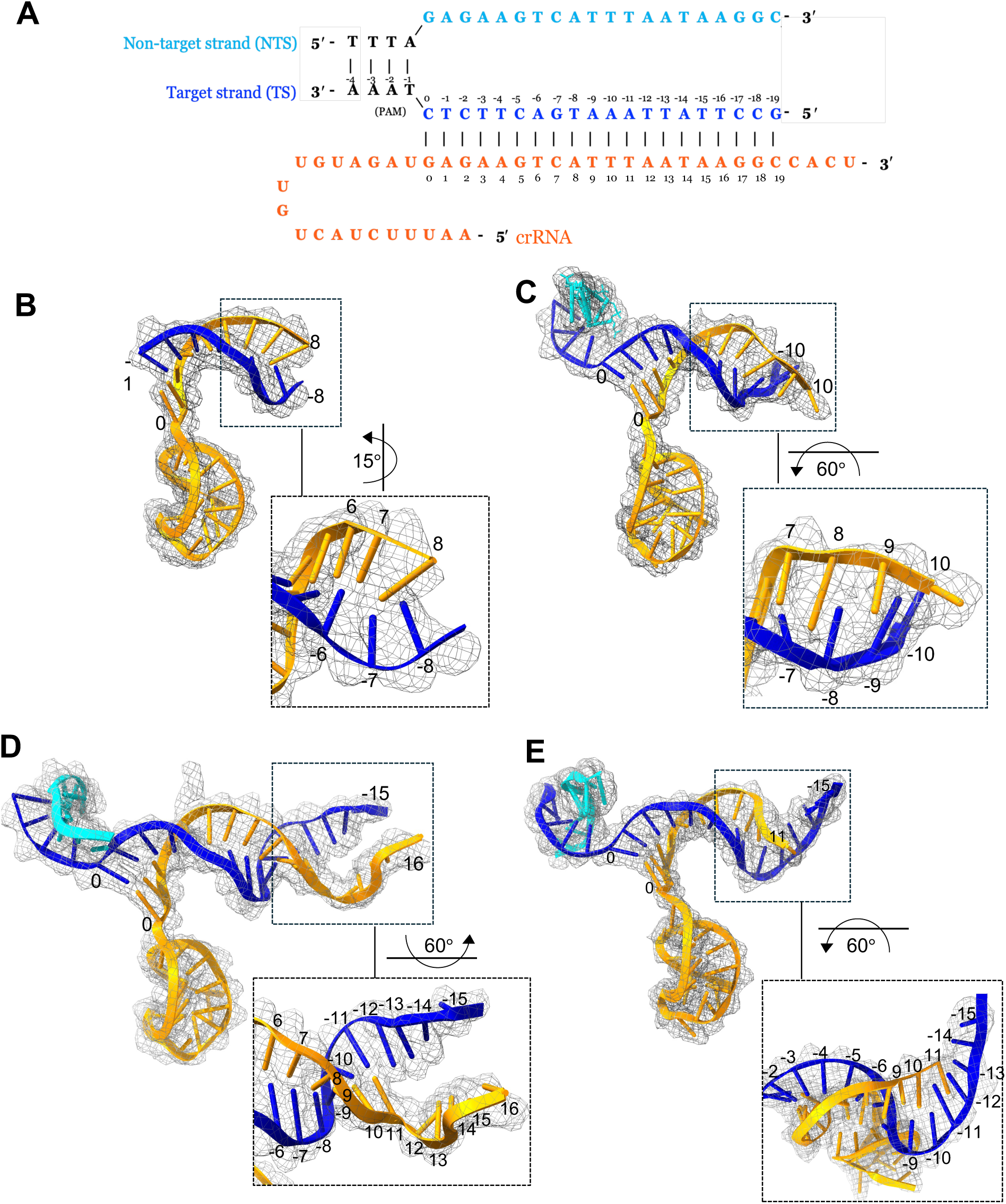
R-loop propagation in the different states of FnoCas12a^KD2P^. **(A)** Schematic showing the base pairing pattern in the R-loop. PAM is in black, and crRNA is in orange. **(B-F)** Map to model fit of R-loop propagation as observed in states S2 **(B)**, S3 **(C)**, S4a **(D)**, and S4b **(E)**. Two different orientations are shown to see the base pairing pattern at different regions of the R-loop (also see **Supplementary Table 6**).

### Helix-1 from RuvC domain is adopted for coordinated conformational changes with the BH

The coordinated conformational changes of BH with the first helix of the RuvC domain (helix-1) was previously proposed^18^ and its structural mechanisms are revelated in FnoCas12a^KD2P^ structures. To further understand the role of the BH and helix-1 in Cas family proteins, we compared the sequences and structures of BH and helix-1 from several Cas12 subtypes and ancestors including TnpB and other DNA enzymes such as resolvase and transposase. A search of the HMMER^38^ database with BH and helix-1 sequences of Cas12a resulted in hits from only Cas12a subtype indicating that the sequence conservation around this region is limited (Supplementary Fig. 12).

We next analyzed whether there is structural conservation of these two helices in Cas12 subtypes and ancestral proteins. Structural similarity search using BH and RuvC domain (FnoCas12a BH and RuvC motif; PDB: 5NFV^25^) as the query identified similar structural folds in other Cas12 subtypes, deoxyribonuclease RuvC, resolvase, and TnpB. This is consistent with previous studies that showed structural similarity between TnpB and Cas12 protein family^39^ and TnpB as the ancestor for Cas12 proteins^40^. The structural similarity dendrogram showed two major clades with the endo-deoxyribonuclease occupying its own clade while the rest of the structures grouped into the other major clade that again branched into 4 sub-clades (Fig. 5A). These observations show that there are some inherent structural differences between the Cas12 and ancestor proteins, which may translate to differences in conformational activation processes in these proteins.

**Figure 5:**
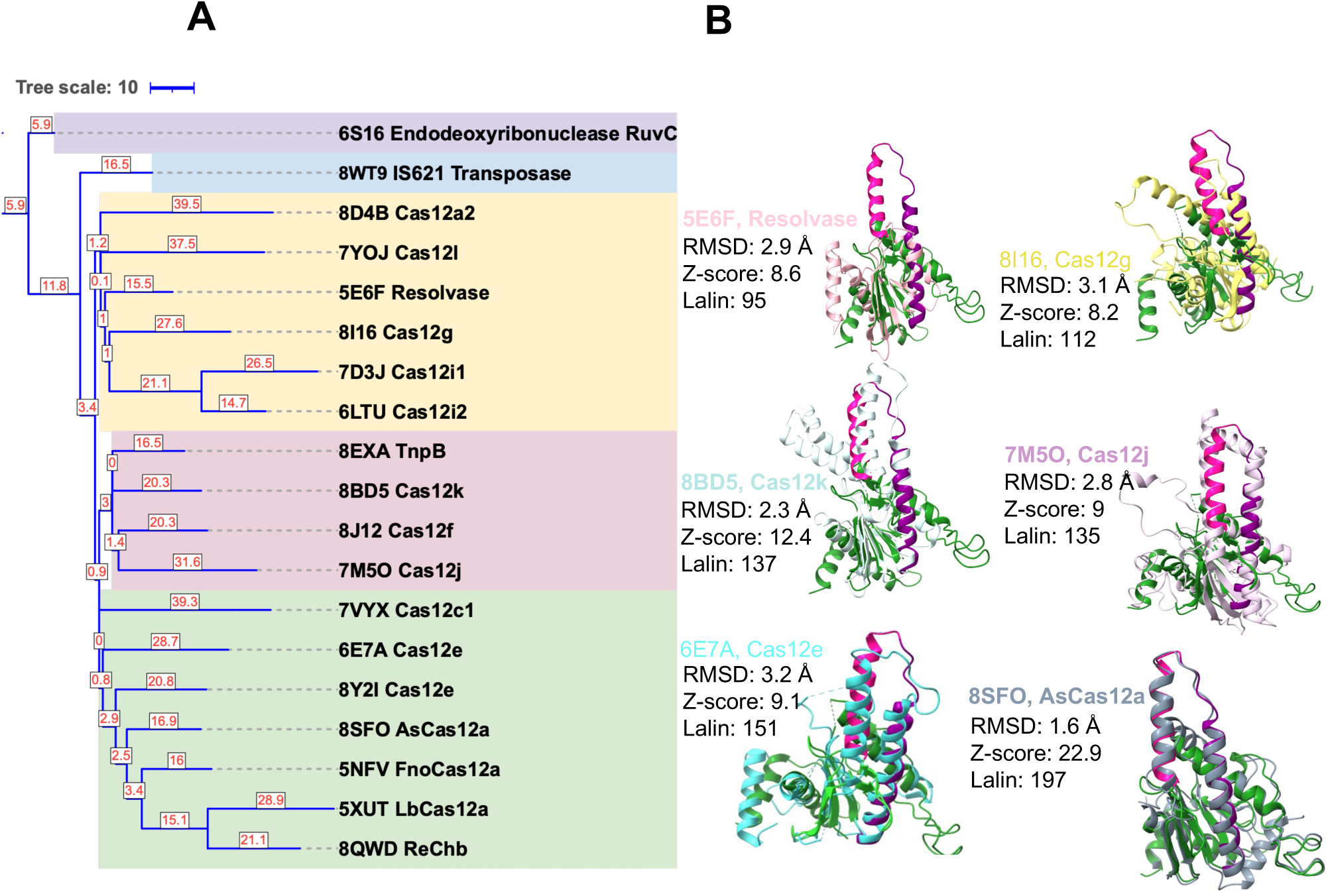
Analysis of the BH and helix-1 conformations of different Cas12 families. **(A)** A DALI Dendrogram representing the hierarchical clustering of structures based on the matrix of DALI z score. A structural homology search for the BH and RuvC domain (∼200 amino acids) of FnoCas12a (PDB: 5NFV) against PDB displayed structural similarity with other Cas12 subtypes and ancestors. The top hits belong to Cas12a orthologs with z scores ranging from 24.9 to 13.9, while hits belonging to other Cas12 subtypes, resolvase, deoxyribonuclease RuvC, and TnpB had z scores ranging from 13.5 to 8.0 indicating there is probable structural homology^40^. iTOL tree^63^ in a rectangle mode, rooted to endodeoxyribonuclease RuvC, was used for visualization and annotation. PDB IDs with the corresponding protein are indicated at the tip of each respective leaf. **(B)** Structural superposition of FnoCas12a (BH in deep pink, helix-1 in purple and the remaining part of the RuvC domain in green) with few representative Cas12 subtypes and other proteins belonging to a subclade. The z score, RMSD, and number of equivalent residues aligned (Lalin) based on the DALI search are shown for each superposition. Each structure aligned against FnoCas12a is shown in one color that matches the color of the PDB ID label associated with it.

To further understand the structural similarities of BH and RuvC motif-II of FnoCas12a, we analyzed the structural overlays of the hits from the DALI (Distance Matrix Alignment) server^41^ (Fig. 5B and Supplementary Fig. 13A). There is a similar organization of BH and helix-1 between FnoCas12a, Cas12f^42^, Cas12j^43,44^, Cas12k^45^, Cas12g^46^,Cas12c1^47^, and Cas12i2^48^. The relative orientation of the two helices is similar in these even though in some cases the connection between the two helices is longer compared to only a loop in between the helices in FnoCas12a. In certain proteins while the BH is in a nearby position without overlaying with FnoCas12a’s BH (e.g., Cas12e^49^, Cas12a2^50^ with an insertion in place of BH, and Cas12l^51^ with BH folded into two helices), the helix-1 still overlays well with that of FnoCas12a (Supplementary Fig. 13A).

Overall, these analyses indicate that helix-1 from RuvC motif-II is co-opted by Cas12 families to work in coordination with the BH to orchestrate conformational changes needed for its activity. Cas9 may have a different mechanism solely focused on its elongated BH for conformational activation since the RuvC motif-II is located farther from the BH in the primary structure (Supplementary Fig. 13B). Interestingly, in the apo^52^ structure, the BH seems to be placed closer to a short helix, located at the start of RuvC-II motif, which gets displaced farther in the binary^53^ and ternary^12^ structures (Supplementary Fig. 13B). Further analysis of multiple Cas proteins will enable deeper characterization of BH-mediated protein activation mechanisms.

## DISCUSSION

A previously reported FnoCas12a^WT^ cryoEM study and our present work on FnoCas12a^KD2P^ have captured several intermediate states through which FnoCas12a-crRNA complex navigates towards attaining the activated pre-catalytic state. The new results from FnoCas12a^KD2P^ shows that an intact BH that can undergo a stable loop-to-helical transition along with R-loop propagation is important in attaining the pre-catalytic stage. It also points to slowing down of the conformational cascade, with more intermediate states that are captured between I2 and I3. We see one state with complete absence of REC2 domain, which was also observed in AsCas12a in earlier stages of R-loop formation and shown to be important in connecting the active state conformation to full R-loop formation. In addition, the elongation of the BH due to the addition of the new helical turns and its bending towards the RNA-DNA hybrid appear to be critical in providing an anchorage for R-loop and to complete the RNA-DNA hybrid formation. In FnoCas12a^WT^ structure, states I2 and I3 have BH in a state prior to the helical conversion. In these structures, BH is at a distance of 17 Å from the RNA-DNA hybrid and a total of 8 base pairs is visible between the crRNA-guide and target DNA. In comparison, in state I4, we see fully formed and bent BH that is at 6 Å to the RNA-DNA hybrid (Supplementary Tables 7 and 8). In FnoCas12a^KD2P^, which is not capable to undergo the loop-to-helical transition, we only see up to a maximum of 11 nt of base pairing between the crRNA-guide and target DNA, with a distance of ∼20 Å from the BH. Interestingly, we also see stacking and distortion of bases towards the end of the R-loop implicating that the strands are not aligned well with an impaired BH, hinting at the essentiality of the stability provided by the BH in these intermediary stages to drive R-loop propagation. The slowing down of the structural transitions is supported by the lower reaction kinetics of FnoCas12a^KD2P^ when compared to FnoCas12a^WT14^.

Opening of the lid of the RuvC active site pocket enables passage of DNA into the active site. In all the structures of FnoCas12a^KD2P^ ternary complex, we observed that the lid is in a loop form occluding the RuvC active site. We propose this to be an allosteric control to prevent non-specific DNA cleavage. The BH-allostery can potentially time the lid opening to only when full complementarity is present in the RNA-DNA hybrid. In another role, the loop-to-helical transition of RuvC’s lid has been shown to play an important role in bringing the TS strand into the active site pocket for DNA cleavage^23,30^. While we do not have structural features to support the TS entry mechanism due to the presence of partial DNA strands, BH’s helical transition most probably contributes towards this second strand passage and cleavage as well. One of the supporting experimental evidences for the bigger contribution of the BH in TS cleavage comes from the activity assays with FnoCas12a^KD2P14^. There is a 6-fold reduction in TS compared to only a 3-fold reduction in NTS cleavage by FnoCas12a^KD2P^ compared to that by FnoCas12a^WT14^. Additionally, FnoCas12a^KD2P^ discriminates the RNA-DNA mismatches by preventing TS cleavage since it was able to nick substrates with mismatches but not linearize them^14^. The AsCas12a structures probing the TS strand entry mechanisms (PDB: 8SFP, 8SFQ and 8SFR^23^) also support these since the BH is extended towards the RNA-DNA hybrid in these structures.

We also observed that FnoCas12a^KD2P^ is unable to perform *trans-*cleavage^14^. The *trans*-cleavage occurs through keeping the RuvC domain “open” for non-specific ssDNA to access the active site^24,54^. The impairment of the BH’s helical transition likely blocks the prolonged activated form of the RuvC pocket after the PAM-distal DNA piece leaves the active site following *cis* cleavage, preventing *trans*-cleavage in FnoCas12a^KD2P^. Similarly, with mismatched DNA, the added energetic cost of unpaired RNA-DNA along with the impaired BH, prevents TS cleavage in FnoCas12a^KD2P^. Further studies are needed for finding the mismatch sensing mechanism of FnoCas12a^KD2P^, especially to understand why positions 12-18 of the RNA-DNA paired region is more sensitive compared to the rest of the R-loop positions^14,55,56^.

Cas12a goes from a closed to open to slightly closed conformation during the different steps following DNA binding and R-loop propagation (Fig. 6)^23^. Our results and previous literature on FnoCas12a^WT^ show that the unwinding of the RuvC helix-1 is a prerequisite to the formation of new turns in the BH. We can envision the helix melting and helix formation as a method to control the degree to which the crab-claw-shaped protein will open the REC2 and Nuc domains to control DNA access in the middle of the bilobed structure. The BH and helix-1 appears to be the two ends of the anchor needed for coordinating the conformational transitions by changing the number of turns in both the helices. The proposed mechanism is supported by previous work that showed complete absence of Cas12a’s DNA cleavage when helix-1 is deleted, and partial activity when BH is deleted^18^. Comparison of the BH conformational changes with AsCas12a^23^ and LbCas12a^27,29,34^ also show similar features. While the BH is elongated in AsCas12a even with only 5 nt base pairing of the crRNA guide and the target DNA, the bending of the BH to reach the RNA-DNA hybrid only happens after 15 nt of base pairing (Supplementary Fig. 14), supporting our proposal that BH’s conformation change is a trigger to support full R-loop formation, which then translates to opening of the RuvC lid^23^. Unwinding of helix-1 is also noticed only in the later stage of R-loop formation (Supplementary Fig. 14), supporting a common mechanism for Cas12a’s DNA cleavage in the orthologs. Our structural comparison of the BH and helix-1 of the diverse Cas12 subtype proteins also hint towards the conservation of these two helices, even in subtypes where there is a domain insertion in between these helices (e.g., Cas12j^43,49^). Interestingly, Cas9 BH is longer and does not show a coordinated helical transition with a RuvC helix. Further studies targeting these helices of Cas9 and Cas12a will help elucidate the differences and similarities of the BH-mediated conformational activation and DNA mismatch sensing mechanisms of these two proteins that evolved differently to perform the same RNA-guided DNA cleavage activity.

**Figure 6:**
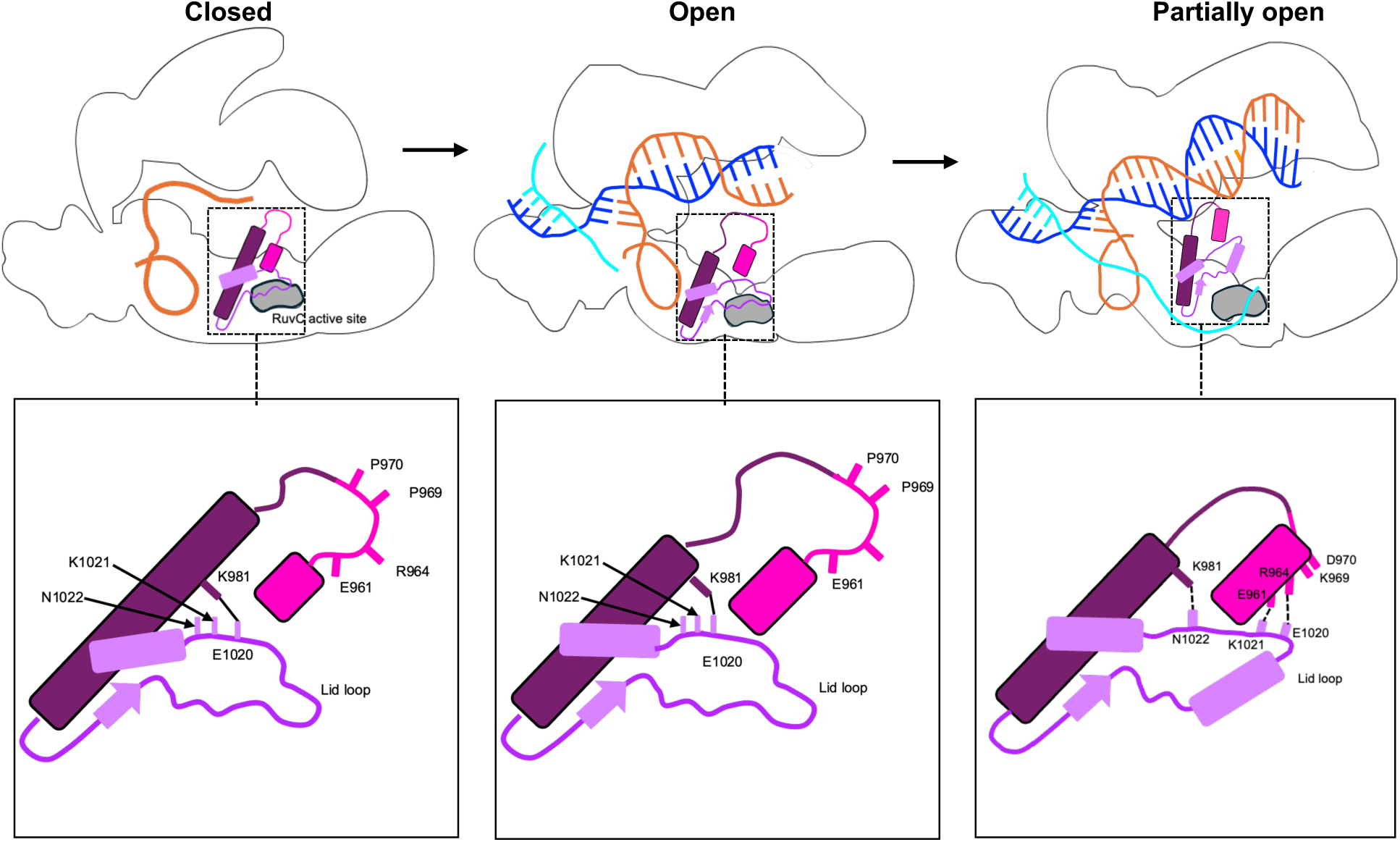
A model showing the central role that BH plays in coordinating the conformational activation of Cas12a. Cas12a undergoes largescale conformational changes that makes the bilobed protein to take a closed conformation after binding to crRNA, followed by an open conformation to enable R-loop growth, and finally attaining a slightly-less opened state after full R-loop formation to reach the pre-catalytic state. Our work illuminates the molecular mechanisms by which BH orchestrates these conformational changes. In the binary complex, part of the BH is shorter with a disordered loop. After DNA binding (∼10 bp of RNA-DNA hybrid formation, Supplementary Fig. 14), the long helix-1 of RuvC domain unwinds a few turns and the BH remains shorter with the loop region. This simultaneous helicity change may enable opening of the bilobed structure. When the R-loop attains ∼16 bp hybridization, the BH elongate by gaining a few helical turns with its concomitant bending towards the RNA:DNA hybrid. The helical change of the BH also helps to open up the RuvC active site pocket by assisting the loop-to-helical transition of the RuvC lid through direct interactions between amino acids in the region of the BH that has undergone the helical transition. Helix-1 acts a pivotal point by changing the residue that it interacts with in the lid before and after its helical transition. In our variant, FnoCas12a^KD2P^, where BH transition is hindered, the protein attains the intermediate conformation (second state in the top panels) but does not proceed to the pre-catalytic state due to its inability to support attainment of the full R-loop in the absence of the loop-to-helical transition of the BH. The lid also exits in a loop form in all our structures. Thus, our structures demonstrate the cooperativity of BH and helix-1 in allosterically opening the RuvC active site pocket based on the number of base pairs in the RNA-DNA hybrid.

In conclusion, cryoEM studies of our variant, FnoCas12a^KD2P^, unravel five intermediary steps that broaden our understanding of the critical role played by BH towards attaining the pre-catalytic state. The structures show that impairment of BH through proline substitutions arrest the conformational cascade, illuminating the allosteric role that BH exerts on the RuvC catalytic pocket by keeping the length of the RNA-DNA hybrid as a checkpoint. The structures also show direct evidence on the cooperativity of BH and helix-1 in enabling R-loop propagation and opening of the RuvC lid. BH being a conserved single helical domain in Cas9 and Cas12 families, as well in other RNA-binding proteins such as the RNA Polymerase, our study reveals the molecular mechanisms by which BH conformational changes regulate communication in multi-domain proteins to assist enzyme function.

## MATERIALS AND METHODS

### Protein expression and purification

The plasmid containing the Fno *cas12a* gene with the K969P/D970P substitutions was purified using previously established procedures^14,57^. *Escherichia coli* Rosetta (DE3) cells containing the FnoCas12a K969P/D970P expression plasmid were grown in 2x Yeast Extract Tryptone (2xYT) and induced with 0.2 mM IPTG at an optical density (OD)_600_ of 0.6-0.9, followed by an overnight incubation at 18 °C. The recombinant protein was purified using a two-step method comprising Ni^2+^-NTA (nitrilotriacetic acid) and cation exchange chromatographies. The purified protein was concentrated using the Thermo Scientific Pierce^TM^ protein concentrator PES, 2-6 mL, 30 kDa and the Cytiva Vivaspin 500 mL, 10 kDa centrifugal protein concentrators and stored on ice until complex preparation for cryoEM.

### RNA and DNA preparation (for oligo sequences see Table S11)

The target strand (TS) and non-target strand (NTS) for the dsDNA target (Fig. 4A) were ordered as ssDNA oligos from SigmaAldrich. The TS and NTS were annealed together by combining equal micromolar amounts of each strand with annealing buffer (10 mM HEPES, pH 8, 50 mM KCl) to a final concentration of 400 μM, followed by heating to 95 °C for 5 min and allowing it to slowly cool to room temperature.

RNA was prepared by *in vitro* transcription and purified as previously described^14,57^. Briefly, template and T7 promoter DNA oligos were ordered from Integrated DNA Technologies (IDT) and annealed in annealing buffer in a 1:1.5 molar ratio of template:T7 promoter strands. The annealed template was used for *in vitro* transcription using T7 RNA polymerase as previously described^57^. The transcription reaction was treated with DNase I to remove the DNA template, and then the crRNA product was precipitated through ethanol precipitation and further purified using a 12% polyacrylamide-8M urea gel. The purified crRNA was suspended in annealing buffer and heated to 95 °C for 2 min then slowly cooled to room temperature for correct folding and quantification. The folded crRNA was stored at 4 °C until use.

### CryoEM grid preparation and data collection

Freshly purified 10 μM FnoCas12a^KD2P^ protein and 12 μM crRNA were incubated at 37 °C for 30 min to form the ribonucleoprotein (RNP) complex in the complexing buffer (20 mM HEPES, pH 7.5, 150 mM KCl, 1% glycerol, 1 mM TCEP, 2 mM EDTA). For formation of the ternary complex, 15 μM DNA duplex was added to the assembled RNP and incubated at 37 °C for another 30 min. The resulting ternary complex was loaded onto a Superdex 200 increase 10/300 GL column (GE healthcare) which was pre-equilibrated with 1X complexing buffer at 4 °C, to separate complexes from excess nucleic acids. Peak fractions were analyzed on 6% native gel (Supplementary Fig. 1) and concentration was determined using absorbance at 280 nm as measured on a Nanodrop 8000 Spectrophotometer (Thermo Scientific) considering 1 Abs = 1 mg/mL. Peak fraction (∼0.147 mg/mL, peak 2) containing the band for ternary complex was vitrified on the same day of purification.

The peak fraction was vitrified using Leica EM GP2 plunger at 4°C and 98% humidity. 4 μL of sample was applied to graphene oxide (Sigma-Aldrich, 0.2 mg/mL) coated^35,36^ Quantifoil R1.2/1.3 300 mesh Cu grid, which were glow discharged using PELCO easiGlow^TM^ at 15 mA for 30 s. After sample application, the grid was immediately blotted for 2 s, followed by plunge freezing.

7232 micrographs were collected at the Stanford-SLAC CryoEM Center (S^2^C^2^), an NIH National CryoEM facility on TEMGamma, a Titan Krios G3i (Thermo Fisher Scientific) cryogenic electron microscope operating at 300 keV and equipped with an X-FEG electron source, fringe-free optics, a BioQuantum (Gatan) energy filter and a K3 (Gatan) direct electron detector. Movies were recorded using a 100 μm objective aperture, an energy filter slit of 20 eV, and an electron dose of 1.25 e^−^/Å^2^ per frame at a magnification of 105k with a corresponding pixel size of 0.86 Å for a total exposure of 50 e^−^/Å^2^ (40 frames over 2.13 s, 0.053 s per frame).

### Single particle cryoEM data processing and model building

The dataset was processed using the standard workflow on CryoSPARCv4.4.1^58^. After patch motion correction and contrast transfer function (CTF) estimation, 5538 micrographs were selected for further processing. Particle picking, conducted with blob picker, identified 6,576,476 particles across all selected micrographs. Following careful inspection, 5,142,814 particles were extracted with a box size of 280 pixels.

We began 2D classification with 400 classes. After four rounds of iterative 2D classification, we selected 399,609 particles for *ab-initio* reconstruction and homogeneous refinement. To assess conformational heterogeneity, 3D variability analysis was performed with a filter resolution of 4.5 Å. The variability observed in this step guided a subsequent 3D classification, resulting in six different classes (Supplementary Figs. 1 and 2).

An initial model was built using PDB: 6GTG^22^ for each class. The models were first aligned with their respective maps on ChimeraX^59^, followed by manually rebuilding in Coot^60^ and refinement using Phenix.real_space_refine^61,62^. The REC1 and REC2 domain were individually isolated and fitted manually into maps followed by rigid body fitting and real space refinement. The models were iteratively improved based on the validation metrics provided by MolProbity^63^ and Phenix refinement.

For the S4a model, which exhibited stretched density for the REC2 domain, the placement of REC2 based on the corresponding position of the domain in PDB: 6GTG^22^ showed inadequate fit for the map and the model. Since our initial analysis showed that state 4a is closer to I3 (PDB: 6GTE^22^, RMSD: 2.5 Å, Supplementary Table 2), we superposed 6GTE onto S4a. Following this, we isolated the REC2 domain (residues 340-591) from PDB: 5NFV^25^, which has a higher resolution, and aligned it with the REC2 position of PDB: 6GTE, that was superimposed on S4a model. Subsequently, the REC2 domain of S4a was replaced with the newly aligned REC2 domain. A rigid body fit refinement was performed using Phenix, during which the starting and ending residues of REC2, 340 and 591, respectively, were deleted to break the link between the REC2 domain and the rest of the structure to enable flexible domain placement during refinement. The resultant model was inspected, and residues lacking density were deleted. Finally, the model underwent refinement using Phenix.real_space_refine to finalize the structure. The density fit analysis shows poor fit for the REC2 domain compared to the rest of the regions of the protein for state S4a.

### Bioinformatics

Profile HMM (Hidden Markov Model) was built using HMMER^38^ for the BH and RuvC region based on the sequence alignment using Clustal Omega^64^ for 313 Cas12a annotated sequences retrieved from NCBI (National Centre for Bioinformatics Information). The profile was used to search for similar sequences using hmmsearch (a tool in HMMER) with a default e value of 0.01. Structural similarity search for the BH-RuvC region (PDB: 5NFV^25^) was performed using DALI server^41^ followed by an all against all structure comparison for 19 structures that had a z score > 8.0. The structural alignment was used to generate a DALI dendrogram. Structures that belong to similar bacterial species were removed to avoid redundancy in the structure comparison. For visualization and annotation of the tree we used an online tool iTOL^65^ (Interactive Tree of Life). The PDB codes and the corresponding structures used for structural comparison are indicated in Figures 5 and Supplementary Fig.13.

## ACKNOWLEDGMENTS

We thank the OU Protein Production & Purification Core (PPC) facility for protein purification services and instrument support and the Biomolecular Structure Core (BSC)-Norman for cryoEM experimental support. The OU PPC Core and BSC-Norman are supported by an IDeA grant from NIGMS [grant number P30GM145423]. Part of this work was performed at the Stanford-SLAC Cryo-EM Center (S^2^C^2^) and National Center for CryoEM Access and Training (NCCAT), which are supported by the National Institutes of Health Common Fund Transformative High-Resolution Cryo-Electron Microscopy program (U24 GM129541 and U24 GM129539). The content is solely the responsibility of the authors and does not necessarily represent the official views of the National Institutes of Health. The authors would also like to thank S^2^C^2^ members (Dr. Wah Chiu, Dr. Nathan D. Burrows and Dr. Xu Yang) and NCCAT members (Dr Edward Eng, Dr. Christina Zimanyi and Dr. Aaron Owji) for their invaluable support and guidance for establishing cryoEM studies in the Rajan laboratory. We also thank Dr. Martin Lawrence and Luke Findlay for valuable advice on cryoEM troubleshooting. We acknowledge Dr. Fares Najar at the High-Performance Computing Center of the Oklahoma State University for building the HMM profile for the BH and helix-1 region.

## DATA AVAILABILITY

All data are included in the paper and Supplementary Information. The atomic coordinates and cryoEM density maps have been deposited in the Protein Data Bank and Electron Microscopy Data Bank under accession codes XXXX/ EMD-XXXXX (state 1), XXXX / EMD-XXXXX (state 2), XXXX / EMD-XXXXX (state 3), XXXX /EMD-XXXXX (state 4a) and XXXX / XXXXX (state 4b).

## INCLUSION AND ETHICS STATEMENT

CryoEM data was collected at the Stanford-SLAC CryoEM Center (S^2^C^2^). Experiments including protein purification, protein-nucleic acid complex preparation, preparation of grids for cryoEM data collection, data processing and model building were performed at the University of Oklahoma, Department of Chemistry and Biochemistry. We carefully agreed upon researcher contributions and authorship criteria to ensure equality in research collaborations.

## AUTHOR CONTRIBUTIONS

RR conceived the idea. RR, CG, and LM designed the experiments. SA performed the bioinformatics and structural comparisons of BH and helix-1. All authors contributed to cryoEM model building, data analysis, and manuscript writing and editing.

