## Supplementary Information for "Helical transition of the bridge helix of Cas12a is an allosteric regulator of R-loop formation and RuvC activation"

### Table of Contents

Supplementary Figures:

Supplementary Fig. 1: Sample purification for cryoEM

Supplementary Fig. 2: Overview of cryoEM data processing workflow

Supplementary Fig. 3: Resolution analysis for the different states

Supplementary Fig. 4: Analysis of the crRNA and DNA in state S1

Supplementary Fig. 5. Setup to analyze the REC1 and REC2 domain movements

Supplementary Fig. 6: Conformational intermediates of FnoCas12a$^{KD2P}$ complex

Supplementary Fig. 7: Analysis of the domain movements across the whole protein in the different FnoCas12a$^{KD2P}$ states

Supplementary Fig. 8: Comparison of the BH, helix-1, and RNA-DNA hybrid transitions in FnoCas12a$^{KD2P}$ and FnoCas12a$^{WT}$

Supplementary Fig. 9: Comparison of the positioning of LKL region of PI domain and HLH region of REC1 domain in the different conformational states FnoCas12a$^{KD2P}$ and FnoCas12$^{WT}$

Supplementary Fig. 10: Interaction of REC-linker and finger region (purple) with crRNA:TS DNA hybrid

Supplementary Fig. 11: Cooperativity of BH and lid in the different conformational intermediates of FnoCas12a$^{KD2P}$

Supplementary Fig. 12: Sequence conservation analysis of Cas12a BH and helix-1 region

Supplementary Fig. 13: Structural conformation of BH and helix-1 in Cas12a and that in Cas9

Supplementary Fig. 14: Conservation of the loop-to-helix transition and bending of BH with R-loop progression in Cas12a orthologs

<u>Tables:</u>

Supplementary Table 1: CryoEM experimental details and Molprobity validation
Supplementary Table 2: RMSD of FnoCas12a$^{KD2P}$ structures with previously available Cas12a structures
Supplementary Table 3: Movement of REC2 and Nuc domains.
Supplementary Table 4: Distance between the REC1 and REC2 domains between the different states
Supplementary Table 5(A): Movement of the REC1 domain with respect to the NUC lobe
Supplementary Table 5 (B): Movement of the REC2 domain with respect to the NUC lobe.
Supplementary Table 6: Nucleic acid base pairing and visibility in different states
Supplementary Table 7: Bending of BH
Supplementary Table 8: Interaction of BH with RNA:DNA hybrid
Supplementary Table 9: Positional shift of HLH domain
Supplementary Table 10: Movement of REC-linker and finger helix
Supplementary Table 11: Oligonucleotide sequences used for FnoCas12a$^{KD2P}$ complexing

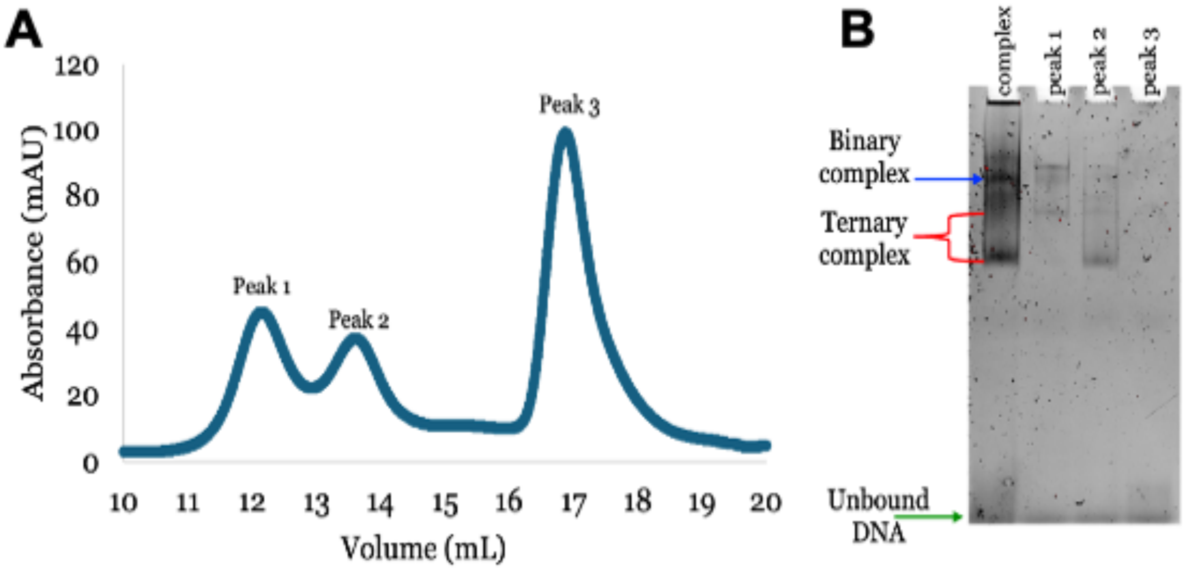

**Supplementary Fig. 1: Sample purification for cryoEM. (A)** A chromatogram showing purification of FnoCas12a$^{KD2P}$ complex using size exclusion chromatography. A molar ratio of 1:1.3:3 of FnoCas12a$^{KD2P}$, crRNA, and DNA was used for the complexing reaction. **(B)** A 6% native gel stained with SYBR green was used to identify the fractions containing the protein-nucleic acid complex. Electrophoretic mobility shift assays had shown compaction of the binary complex upon binding to DNA (data not shown).

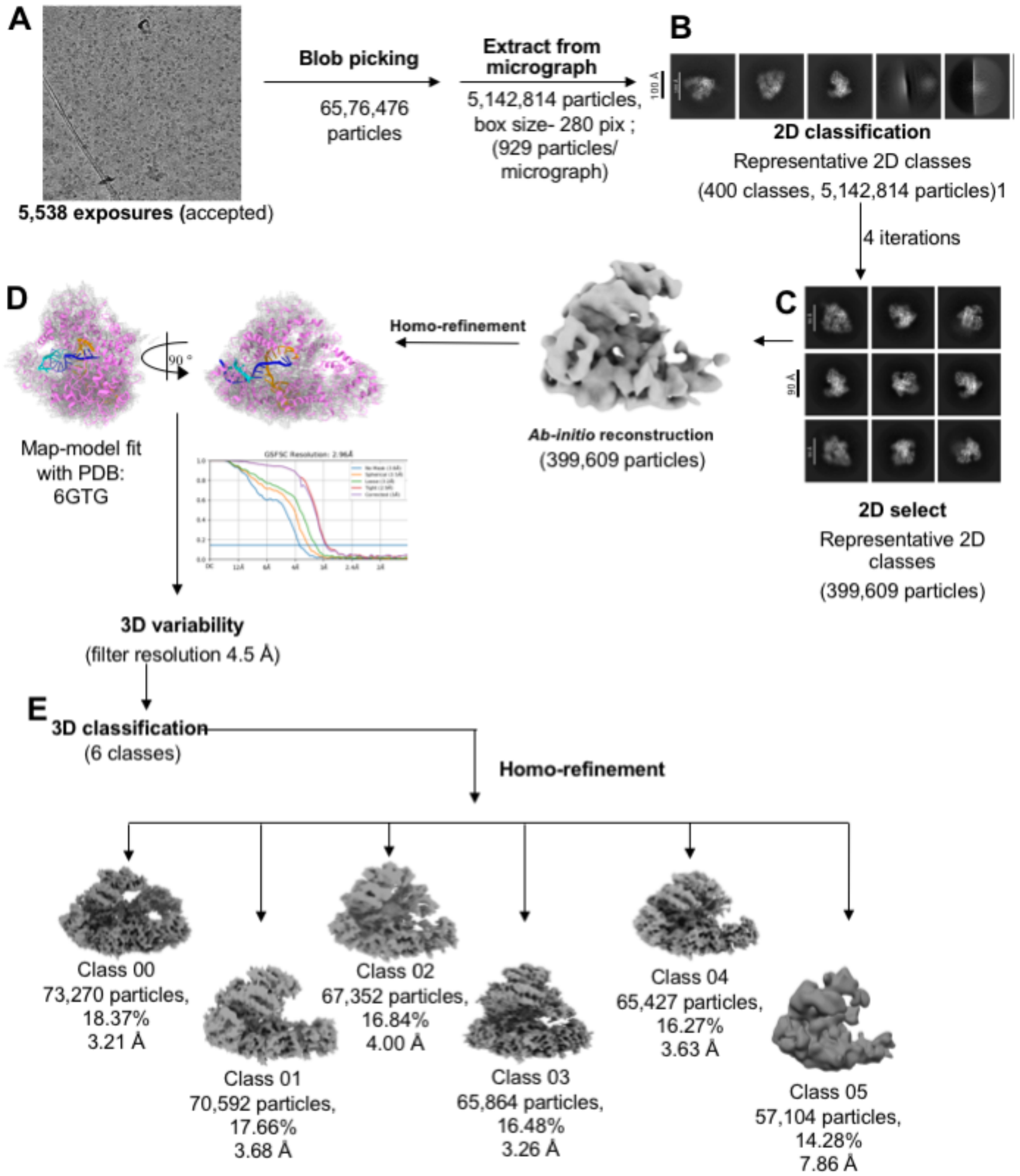

**Supplementary Fig. 2**: **Overview of cryoEM data processing workflow.** Overview of cryoEM data processing workflow. CryoEM data analysis was performed using CryoSPARCv4.4.155. **(A)** Representative cryoEM micrograph of FnoCas12a$^{KD2P}$ complex in vitreous ice on a graphene oxide-coated Quantifoil grid. **(B)** Representative 2D classes from the initial batch of particles. **(C)** Representative 2D classes after performing four rounds of iterative 2D classification to select particles with the expected morphology. **(D)** An *ab-initio* 3D reconstructed volume was obtained and further refined using homogeneous refinement. A model of the FnoCas12a$^{WT}$ structure (PDB: 6GTG) was fit into the homo-refined map. **(E)** 3D variability analysis followed by 3D classification identified six distinct states of FnoCas12a$^{KD2P}$. Models were built for the five classes exhibiting clear features of the Cas12a complex, while Class 05 with a low-resolution was not pursued for model building.

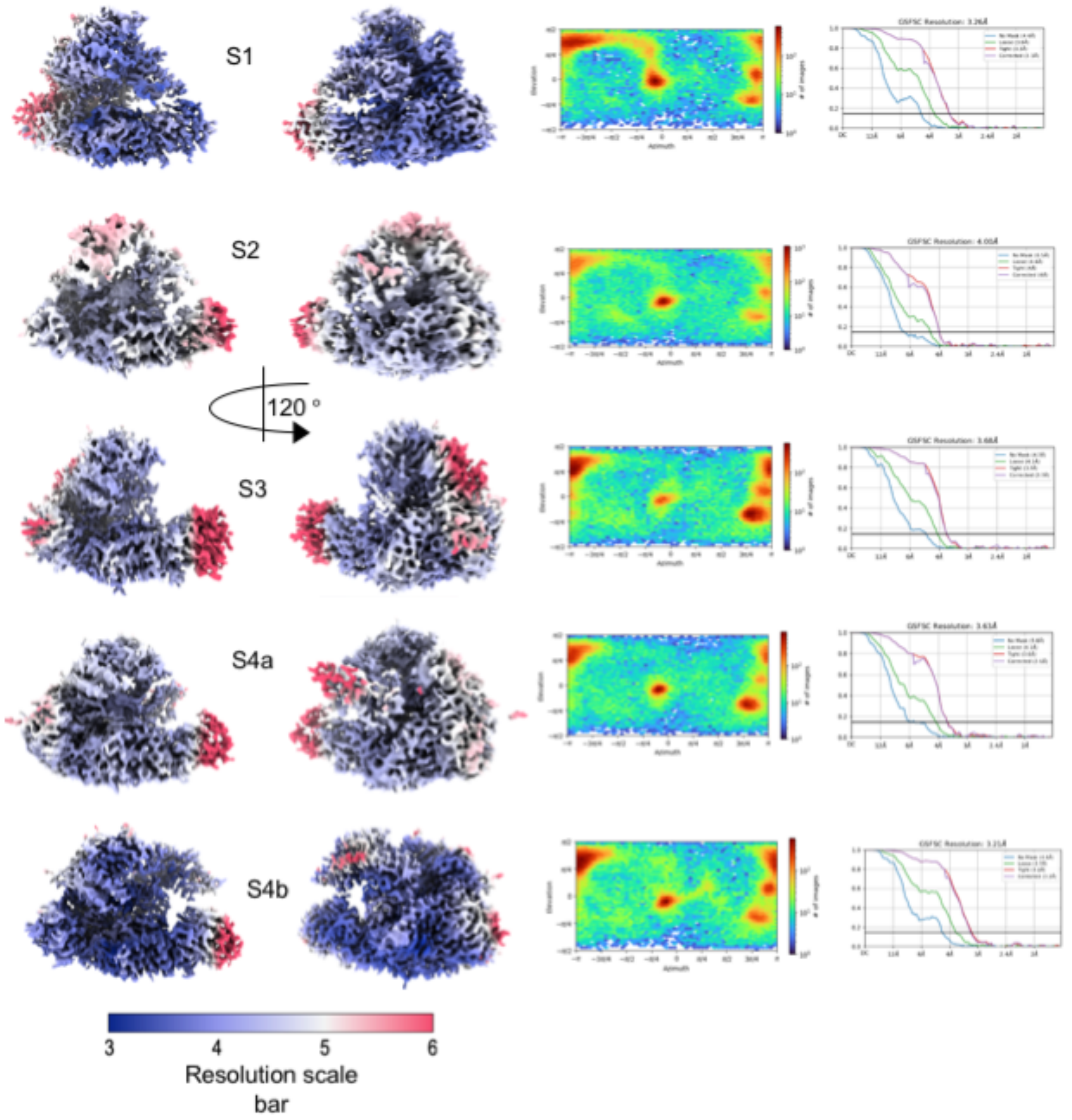

**Supplementary Fig. 3: Resolution analysis for the different states.** The GSFSC resolution for the different states ranges from 3.21 Å to 4 Å (related to Fig.1). For each panel, in left to right order, shown are local resolution maps generated using Phenix, reconstruction Euler diagrams showing particle orientation distributions, and reconstruction gold-standard FSC (Fourier Shell Correlation) curves (resolution estimated at FSC=0.143). Map resolutions and particle distribution plots were produced using CryoSPARC.

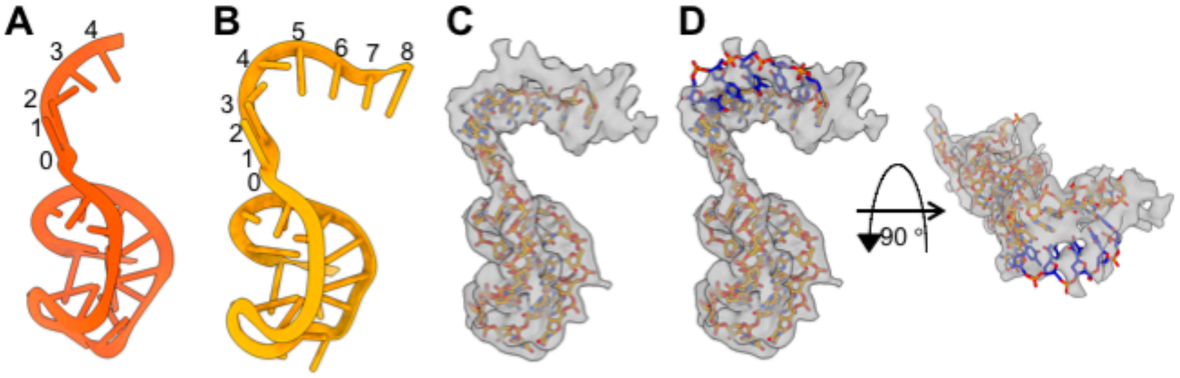

**Supplementary Fig. 4: Analysis of the crRNA and DNA in state S1 (**related to **Fig 1).** Impaired BH does not impact crRNA seed pre-ordering. Comparison of crRNA from S1 **(A)** and FnoCas12a$^{WT}$ binary structure **(B**, PDB: 5NG6**)** shows the ordered seed region for facilitating target DNA search. **(C)** Extracted density for only the nucleic acid in S1 shows residual density at a region that can potentially belong to the TS DNA. **(D)** Superimposing TS DNA (blue) from S2 on S1 map (crRNA, orange belongs to S1) shows extra density in S1 that can hold a TS DNA. This indicates that S1 is a structure at the start of target DNA identification. We did not build TS DNA into this density since it was weak.

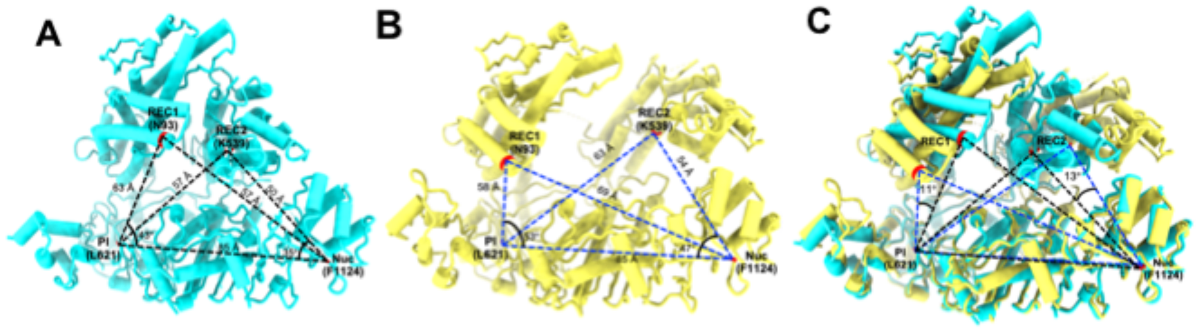

**Supplementary Fig. 5. Setup to analyze the REC1 and REC2 domain movements.** Cas12a model for states S1 **(A)**, S4b **(B)**, and an overlay of both the states **(C)** show the movements of REC1 and REC2 domains in S4b relative to S1 (see also **Tables S5A and S5B**). Residue L621 from the PI domain and F1124 from the Nuc domain, which maintain consistent positions across the different states, were used to define the axis of movement of the REC domains. Residues N93 from the REC1 domain and K539 from the REC2 domain were selected to create two triangles and the angle between the NUC-lobe axis and the REC1/REC2 domains were used to describe the rotation, while distance between residues K539 and F1124 was used to assess the opening of the NUC and REC lobes. Distances were measured using ChimeraX, and angles were calculated using the cosine equation: cos(A) = ((b^2+c^2-a^2))/2bc.

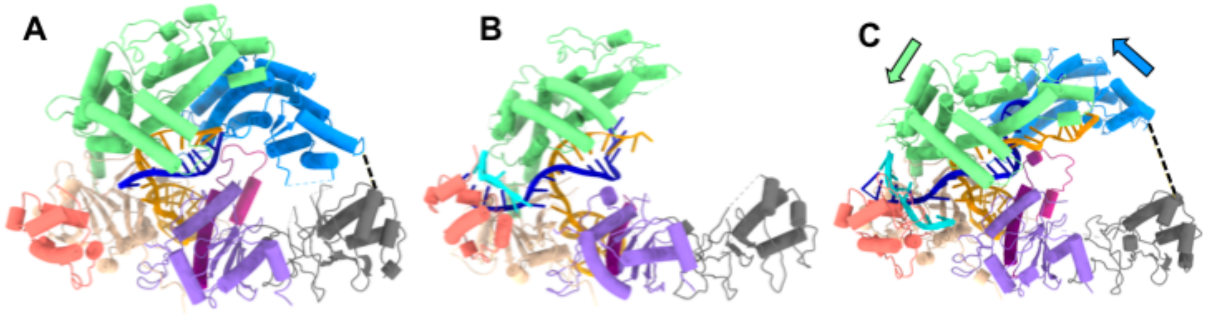

**Supplementary Fig. 6: Conformational intermediates of FnoCas12a$^{KD2P}$ complex (**related to **Fig. 1).** Models built for states S2 **(A)**, S3 **(B)**, and S4a **(C)** with domains colored as in scheme in **Figure 1A**. During the transition from S2 to S4a, the distance between the REC2 and Nuc domain increases by 32 Å and REC1 and REC2 by 24 Å (**Tables S3 and S4**). Arrows depict the direction of the corresponding movements.

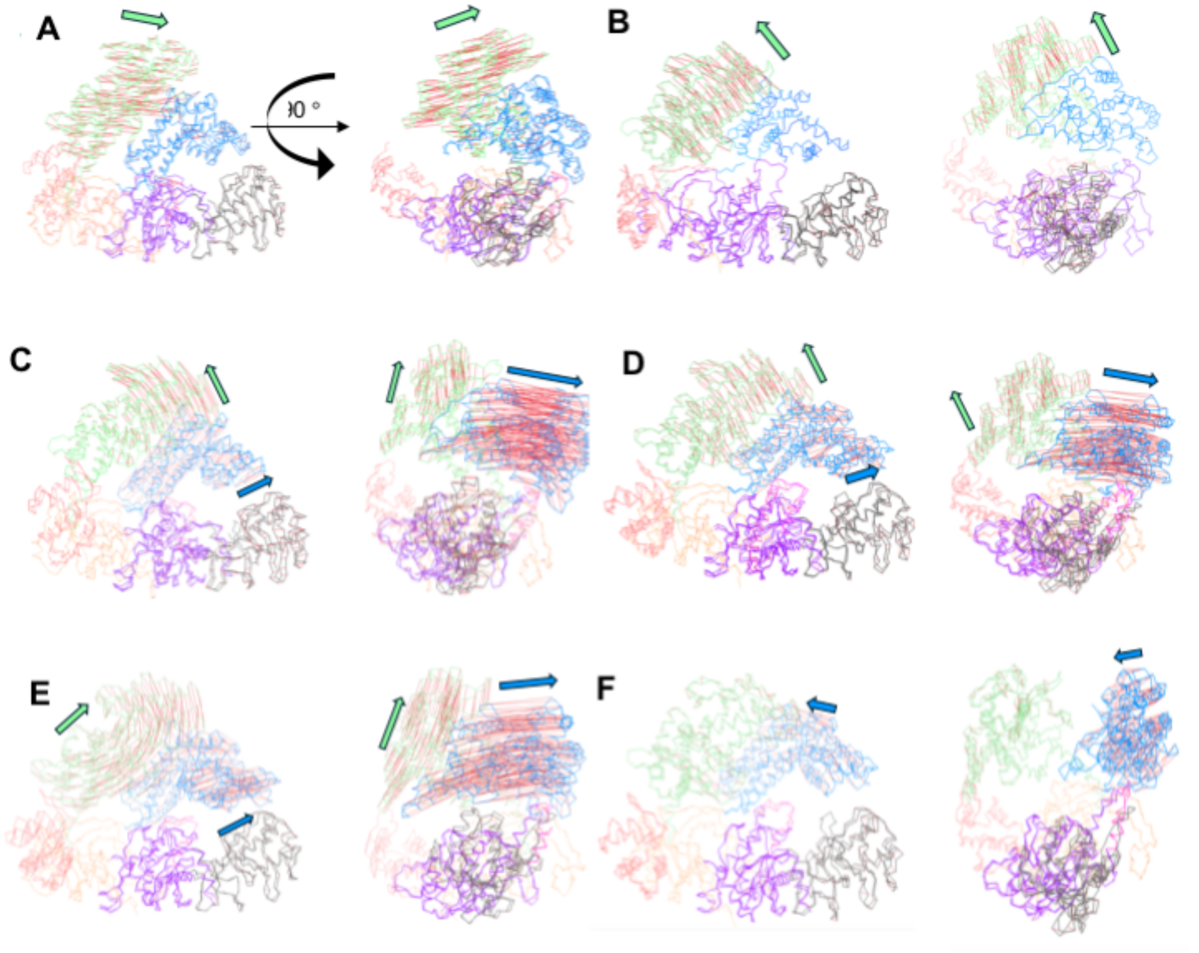

**Supplementary Fig. 7: Analysis of the domain movements across the whole protein in the different FnoCas12a$^{KD2P}$ states.** The red lines represent the magnitude of Ca movement for each amino acid, while the pale green and dodger blue arrows indicate the direction of movement for the REC1 and REC2 domains, respectively, during the transitions between the different states. The vector diagrams were generated using PyMOL[1] (See **Tables S5A and S5B** for translation and rotation values calculated using ChimeraX[2]). **(A)** The REC1 domain shows a significant movement between S1 and S2, moving away from the REC2 domain. **(B)** From S2 to S3, REC1 continues to move away from the central cavity. Comparison between S2 and S4a **(C)** and between S2 and S4b **(D)** reveals significant rotation and translation of the REC2 domain. **(E)** Overall, both REC1 and REC2 domains undergo extensive rotation and translation from S1 to S4b. **(F)** Compared to S4a, the REC2 domain in S4b slightly moves towards the central cavity.

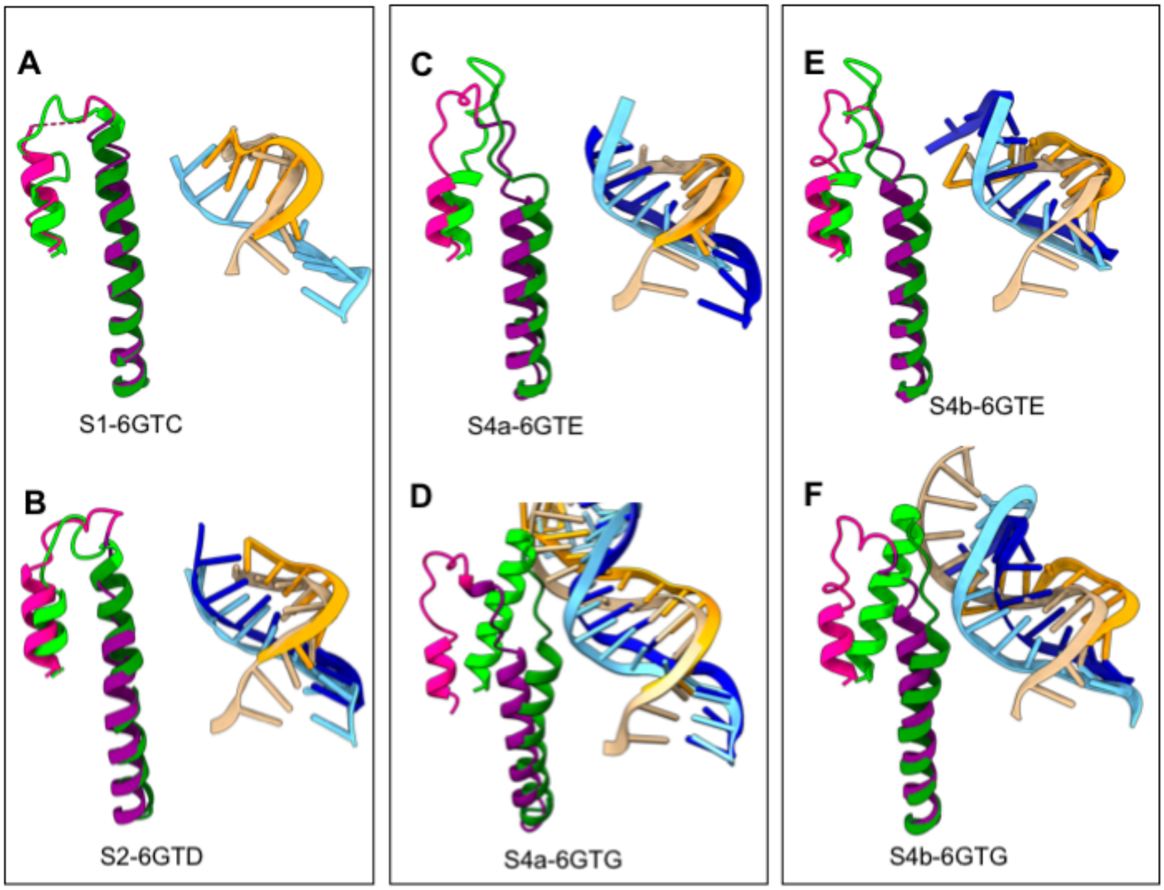

**Supplementary Fig. 8: Comparison of BH, helix-1, and RNA-DNA hybrid transitions in FnoCas12a$^{KD2P}$ and FnoCas12a$^{WT}$** (related to **Fig. 2**). **(A–F)** Structural comparison of BH and helix-1 movements in FnoCas12a$^{WT}$ and FnoCas12a$^{KD2P}$ across the different conformational states. BH is shown in lime green for FnoCas12a$^{WT}$ and in deep pink for FnoCas12a$^{KD2P}$; helix-1 is colored forest green for FnoCas12a$^{WT}$ and purple for FnoCas12a$^{KD2P}$; TS DNA is in sky blue for FnoCas12a$^{WT}$ and medium blue for FnoCas12a$^{KD2P}$; crRNA is in tan for FnoCas12a$^{WT}$ and in orange for FnoCas12a$^{KD2P}$. Comparisons are as follows: **(A)** S1 *vs.* I1 (PDB: 6GTC); **(B)** S2 *vs.* I2 (PDB: 6GTD); **(C)** S4a vs. I3 (PDB: 6GTE); (D) S4a vs. I4 (PDB: 6GTG); (E) S4b vs. I3 (PDB: 6GTE); and (F) S4b *vs.* I4 (PDB: 6GTG). In FnoCas12a$^{WT}$, the BH elongates and bends toward the R-loop. In FnoCas12a$^{KD2P}$ states S4a and S4b, the BH does not undergo loop-to-helix transition or bending and it stays farther away from the R-loop. The unwinding of helix-1 can be seen in FnoCas12a$^{KD2P}$, starting from state S2.

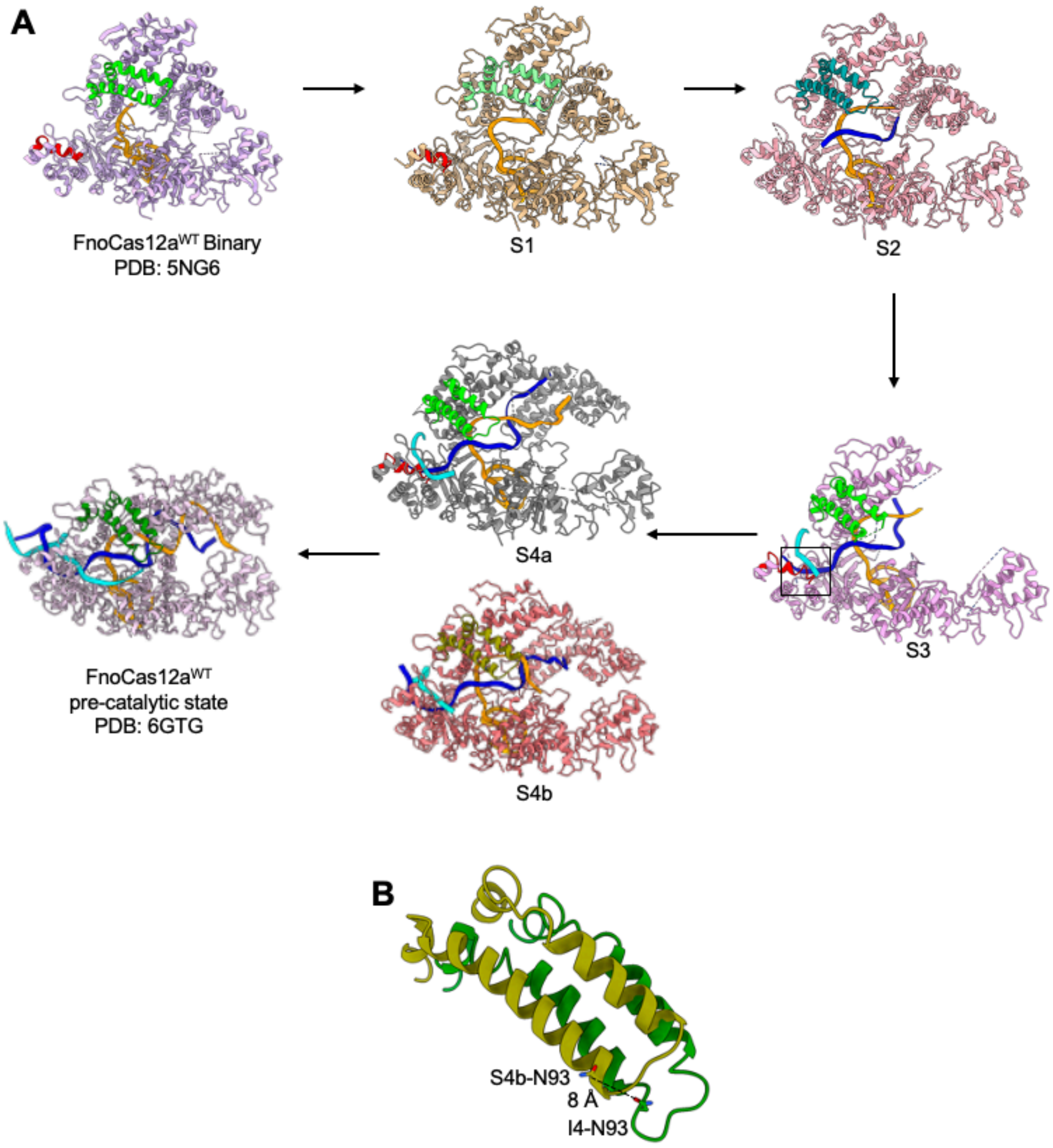

**Supplementary Fig. 9: Comparison of the positioning of LKL region of PI domain and HLH region of REC1 domain in the different conformational states FnoCas12a$^{KD2P}$ and FnoCas12$^{WT}$. (A)** Different conformational states of FnoCas12a$^{KD2P}$ are placed within the catalytic cycle. While the LKL region (red) of the PI domain has similar range of movements as that of FnoCas12a$^{WT}$, movement of HLH region (shades of green) is reduced in the variant. Comparison of HLH segment (S1 in pale green, S2 in teal, S4b in olive, and I4 (PDB: 6GTG) in forest green)

are as follows: **(B)** S2 *vs.* S1; **(C)** S4b *vs.* S2; **(D)** S4b *vs.* I4 (PDB: 6GTG); **(E)** S2 *vs.* I4 and **(F)** S1 *vs.* I4 (PDB:6GTG) (also see **Table S9**).

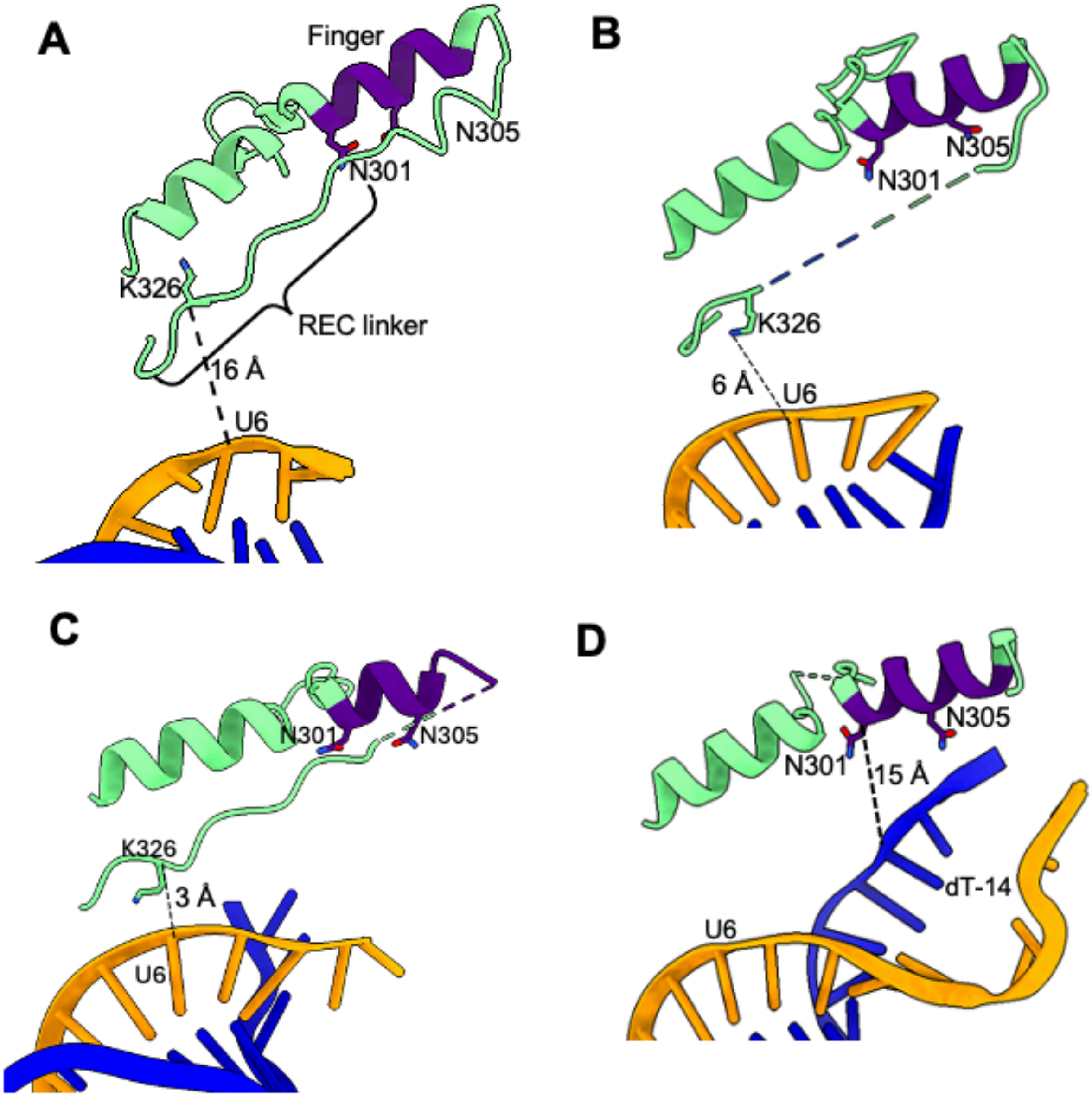

**Supplementary Fig. 10: Interaction of REC-linker and finger region (purple) with crRNA:TS DNA hybrid** (related to **Figure 3**, also see **Table S7**). **(A–D)** Comparison of the REC-linker and finger region movements in 6GTC, S2, S3, and S4a (see **Table S10**). Comparison of all the FnoCas12a$^{KD2P}$ states with the corresponding FnoCas12a$^{WT}$ states shows that the REC-linker attains a similar position in both the proteins. In the FnoCas12a$^{KD2P}$ complex, the finger region remains farther from the PAM distal end of the R-loop compared to the FnoCas12a$^{WT}$ pre-catalytic state (PDB: 6GTG[1]). The distances between the Cα-atoms of the amino acid residues and the

C4′-atom of the nucleotide was measured for analysis. Residue K326 from the REC-linker, and residues N301 from the finger regions, were selected to measure the distances to U6 of crRNA and dT-14 of TS DNA, respectively. The density for REC linker in S4a is missing.

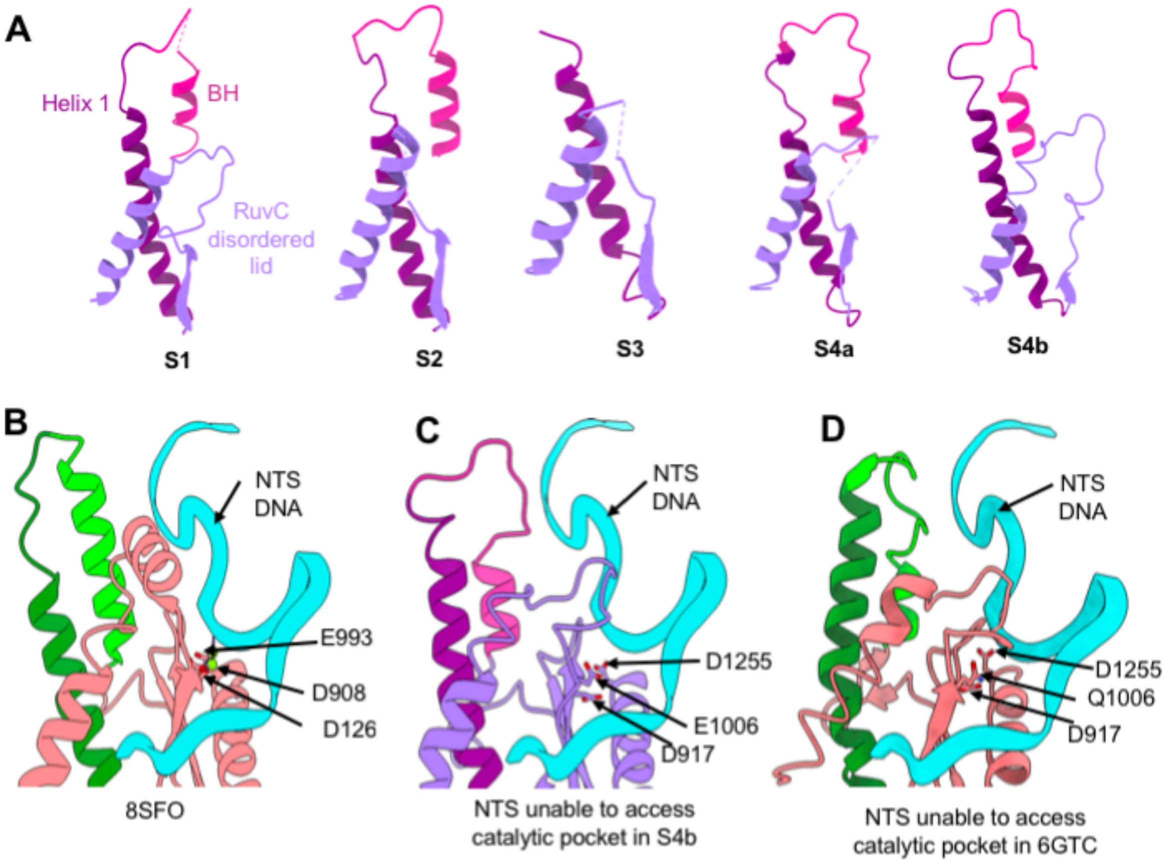

**Supplementary Fig. 11: Cooperativity of the BH and lid in the different conformational intermediates of FnoCas12a$^{KD2P}$. (A)** Overview of the structures of the BH, helix-1, and RuvC lid showing coordination between the different protein components in states S1, S2, S3, S4a and S4b. **(B)** A view of the RuvC catalytic pocket of AsCas12a (PDB: 8SFO). The lid adopts a helical conformation, and the NTS DNA can access the RuvC catalytic pocket. **(C)** An overlay of the NTS DNA from AsCas12a (PDB: 8SFO) into the RuvC catalytic pocket of state S4b of FnoCas12a$^{KD2P}$ and **(D)** an early intermediate of FnoCas12a$^{WT}$ (PDB: 6GTC). The RuvC lid exists in a loop conformation, occluding the catalytic pocket, and preventing NTS DNA entry.

**A**

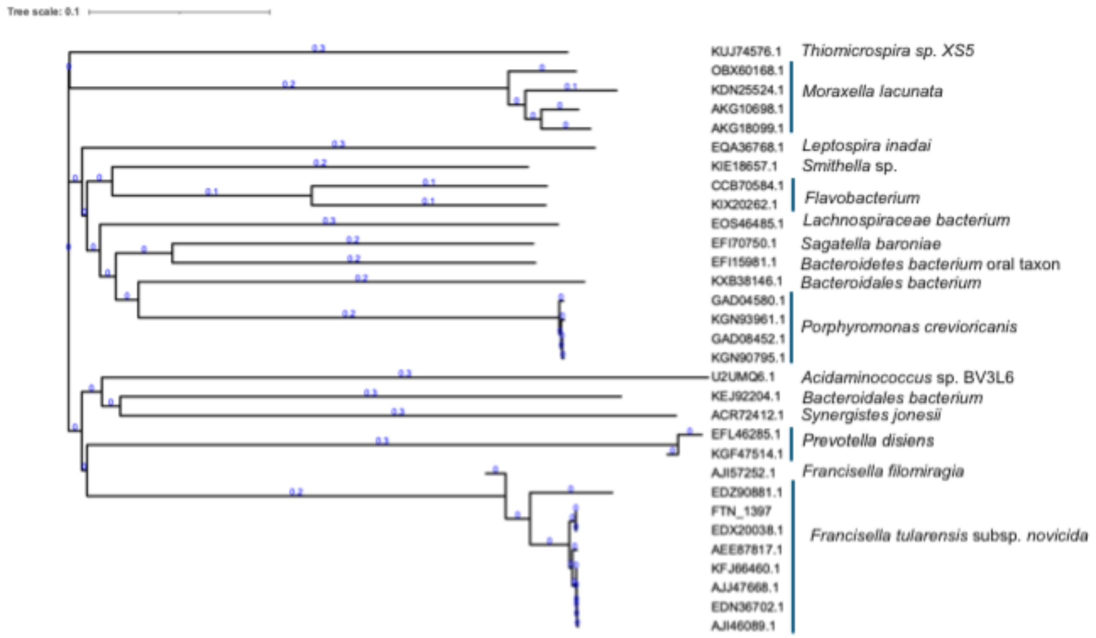

**B**

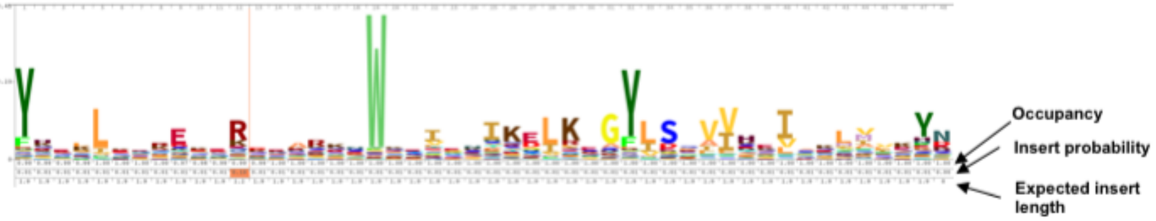

**Supplementary Fig. 12: Sequence conservation analysis of Cas12a BH and helix-1 region. (A)** A dendrogram showing the sequence conservation of BH and helix-1. To generate an HMM profile, Cas12a sequences were aligned using Clustal Omega[2]. Based on the BH and helix-1 region of FnoCas12a, the region encompassing this region in all the other Cas12a sequences were trimmed. An HMM profile for this trimmed region was generated and was used to identify similar sequences using hmmsearch[3]. Thirty-one hits were obtained all of which belong to only Cas12a subtype. The resulting hits were then aligned to build a phylogenetic tree and visualized using iTOL tool[4]. All the hits are grouped under one clade which then branches into subclades. The branch lengths are indicated and the protein ID corresponding to each hit is shown at the tip of each leaf. **(B)** A logo for the HMM profile to represent the conservation of residues in the BH and helix-1 region of Cas12a. The logo was created using the online tool Skylin[5]. The height of the stack represents the conservation of the residue at that position.

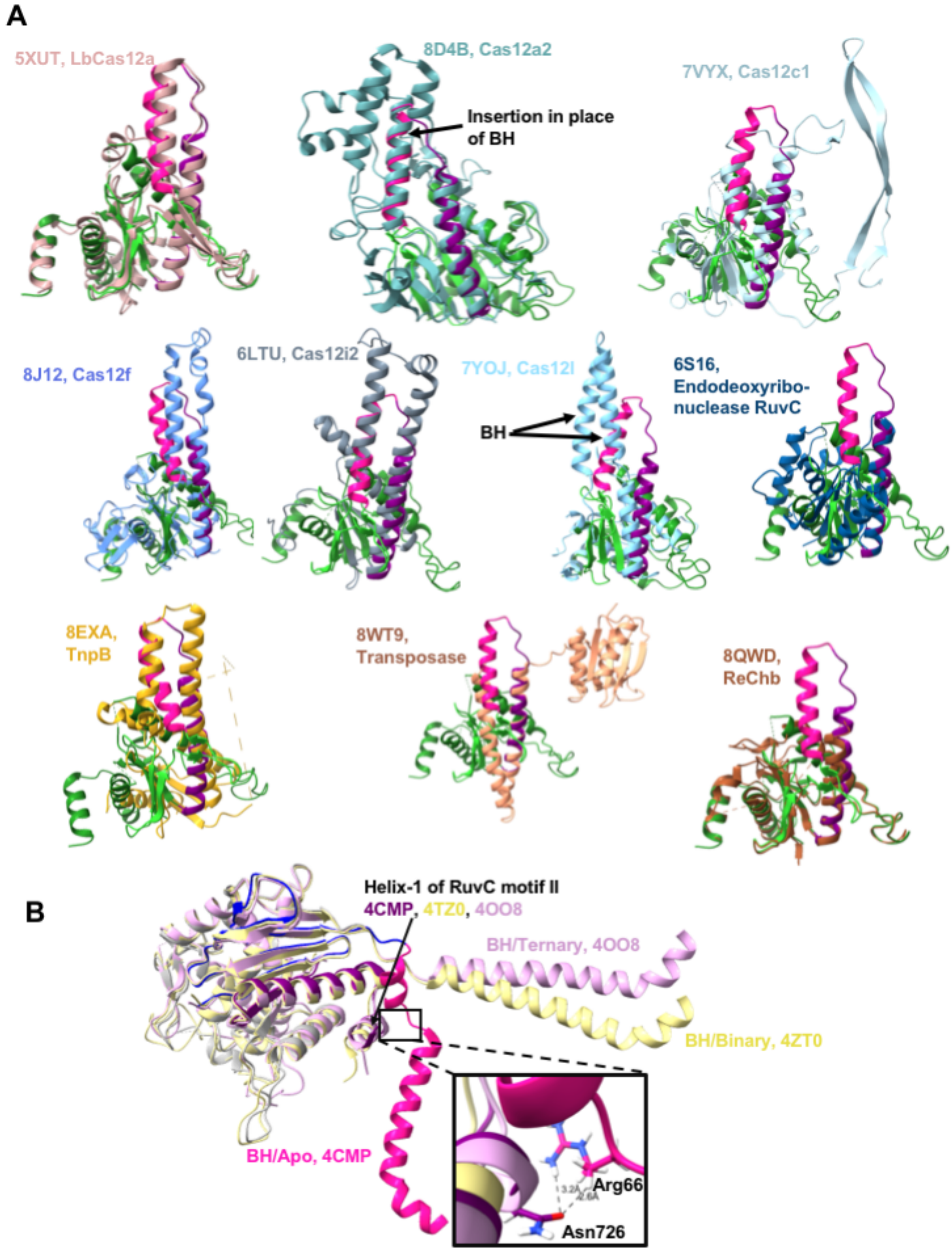
A
5XUT, LbCas12a
8D4B, Cas12a2
Insertion in place of BH
7VYX, Cas12c1
8J12, Cas12f
6LTU, Cas12i2
7YOJ, Cas12l
6S16, Endodeoxyribo-nuclease RuvC
BH
8EXA, TnpB
8WT9, Transposase
8QWD, ReChb
B
Helix-1 of RuvC motif II
4CMP, 4TZ0, 4OO8
BH/Ternary, 4OO8
BH/Binary, 4ZT0
BH/Apo, 4CMP
Arg66
Asn726

**Supplementary Fig. 13: Structural conformation of BH and helix-1 in Cas12a and that in Cas9. (A)** Structural superposition of FnoCas12a (BH in deep pink, helix-1 in purple, and the remaining part of the RuvC domain in green) with Cas12 subtypes and other proteins belonging to a subclade. Each structure aligned against FnoCas12a is shown in one color that matches the color of the PDB ID label associated with it. **(B)** The apo (PDB: 4CMP[6]), binary (4ZT0[7]) and ternary (4OO8[8]) structures of Cas9 from *Streptococcus pyogenes* were superposed to identify similarly coordinating conformations for the BH and helix-1 as in Cas12 subtypes. Unlike that observed in Cas12 subtypes, the BH of Cas9 is flanked by RuvC motif-I on one side and by the REC domain on the other side. The first helix of RuvC motif-II, comparatively shorter in length than that of Cas12a, is placed closer to the BH in the apo structure with a potential interaction between the side chain of Arg66, one of the four residues in the region of the BH that undergoes loop-to-helix transition after binding to the single guide RNA (sgRNA), and Asn726 in the first helix of RuvC motif-II. However, the BH moves farther away in the binary and the ternary structures (~17.4 Å) making no interaction with the first helix of RuvC motif-II, both sequentially and structurally, after Cas9 binds to nucleic acid.

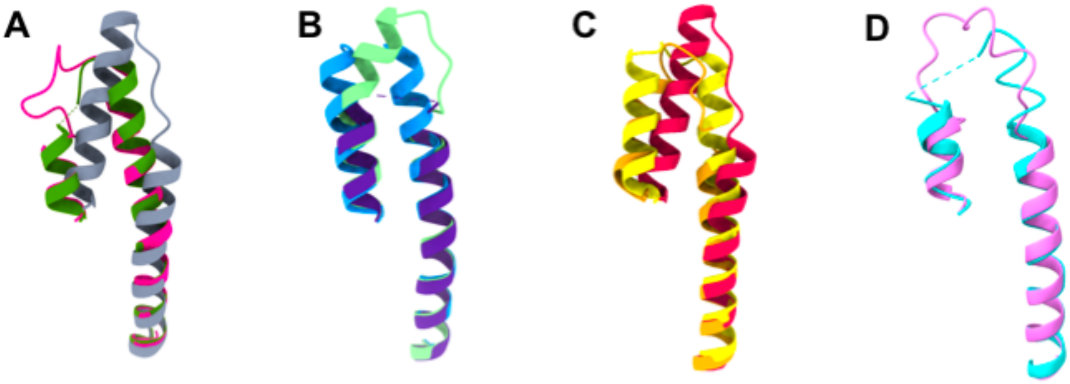

**Supplementary Fig.14: Conservation of the loop-to-helix transition and bending of the BH with R-loop progression in Cas12a orthologs.** In this image, the nucleic acids are not shown for clarity. Comparing the BH of FnoCas12a$^{WT}$ [**A**; binary[9]: forest green; R-loop progression- 8 bp[1]: deep pink; 20 bp[1]: grey], AsCas12a [**B,** R-loop progression- 5 bp: dodger blue; 10 bp: indigo; 15 bp[10]: pale green], and LbCas12a [**C,** binary[11]: yellow; R-loop progression-14 bp[12]: gold, R-loop progression- 20 bp[13], red], the gradual melting of helix-1 followed by bending of the elongated BH towards the R-loop appears to be a conserved mechanism. **(D)** Melting of helix-1 from S1 (cyan) to S4a (pink) is visible, but proline substitutions prevent the elongation and bending of the BH.

**Supplementary Table 1: CryoEM experimental details and Molprobity validation**

| | **S1** | **S2** | **S3** | **S4a** | **S4b** |
|---|---|---|---|---|---|
| | PDB: XXXX | PDB: XXXX | PDB: XXXX | PDB: XXXX | PDB: XXXX |
| | EMD- XXXXX | EMD- XXXXX | EMD- XXXXX | EMD- XXXXX | EMD- XXXXX |
| Data collection and processing | | | | | |
| **Voltage (kV)** | 300 | 300 | 300 | 300 | 300 |
| **Electron exposure (e/ Å$^2$)** | 50 | 50 | 50 | 50 | 50 |
| **Defocus range (μm)** | -1 to -2 | -1 to -2 | -1 to -2 | -1 to -2 | -1 to -2 |
| **Pixel size (Å)** | 0.86 | 0.86 | 0.86 | 0.86 | 0.86 |
| **Symmetry imposed** | C1 | C1 | C1 | C1 | C1 |
| **Initial particle images (no.)** | 51,42,814 | 51,42,814 | 51,42,814 | 51,42,814 | 51,42,814 |
| **Final particle images (no.)** | 65,864 | 67,352 | 70,592 | 65,427 | 73,270 |
| **Map resolution (Å)** | 3.26 | 4 | 3.68 | 3.63 | *3.21* |
| **FSC threshold** | 0.143 | 0.143 | 0.143 | 0.143 | 0.143 |
| Refinement | | | | | |
| **Initial model used (PDB code)** | 6GTG | 6GTG | 6GTG | 6GTG | 6GTG |
| **Map correlation coefficient (main chain)** | 0.74 | 0.74 | 0.77 | 0.74 | 0.78 |
| Model composition | | | | | |
| **Non-hydrogen Atoms** | 10866 | 10735 | 8472 | 11190 | 11756 |
| **Protein Residues** | 1242 | 1207 | 894 | 1189 | 1276 |
| **Nucleotides** | 28 | 36 | 52 | 64 | 58 |
| RMSDs | | | | | |
| **Bond lengths (Å)** | 0.25 | 0.27 | 0.28 | 0.32 | 0.28 |
| **Bond angles (°)** | 0.49 | 0.49 | 0.54 | 0.53 | 0.53 |
| Validation | | | | | |
| **MolProbity score** | 1.79 | 2.02 | 1.82 | 1.89 | 1.82 |
| **Clash score** | 9.53 | 13.81 | 9.09 | 11.42 | 8.99 |
| **Poor rotamers (%)** | 0 | 0.27 | 0 | 0 | 0.09 |
| **Ramachandran plot Favored (%)** | 95.84 | 94.52 | 95.74 | 95 | 95.97 |
| **Allowed (%)** | 4 | 5.4 | 4.15 | 5 | 4.8 |
| **Disallowed (%)** | 0.16 | 0.08 | 0.12 | 0 | 0 |
| **See also Figure S2. FSC, Fourier shell correlation.** | | | | | |

**Supplementary Table 2: RMSD of FnoCas12a$^{KD2P}$ structures with previously available Cas12a structures.** RMSD across all Cα-atoms were measured between the FnoCas12a$^{KD2P}$ states and previously published Cas12a structures (calculated using ChimeraX).

| State | PDB ID | RMSD (Å) |
|---|---|---|
| S1 | 5NG6[9] (FnoCas12a, binary) | 2.4 |
| | 6GTC[1] (FnoCas12a, I1) | 2.6 |
| | 6NME[11](LbCas12a, binary) | 4.6 |
| | 5ID6[14](LbCas12a, binary) | 4.7 |
| S2 | 6GTD [1](FnoCas12a, I2) | 3.6 |
| | 8SFH [10](AsCas12a, 5 bp) | 6.5 |
| | 8SFI [10](AsCas12a, 8 bp) | 1.4 |
| S3 | 6GTD[1] (FnoCas12a, I1) | 3.6 |
| | 8SFJ [1](AsCas12a, 10 bp) | 8.0 |
| | 8I54 [12](LbCas12a, 14 bp) | 9.9 |
| S4a | 6GTD [1](FnoCas12a, I2) | 12.3 |
| | 6GTE[1] (FnoCas12a, I3) | 5.2 |
| | 6GTG[1] (FnoCas12a, I4) | 7.4 |
| | 5NFV[9] (FnoCas12a, 20 bp) | 6.6 |
| | 6I1L [15](FnoCas12a, 20 bp) | 7.5 |
| | 6I1K[15] (FnoCas12a, 20 bp) | 6.6 |
| | 8SFL[10] (AsCas12a,15 bp) | 6.8 |
| | 8SFN [10](AsCas12a,16 bp) | 9.4 |
| | 8SFO [10](AsCas12a, 20 bp) | 10.9 |
| | 8I54[12] (LbCas12a, 14 bp) | 13.9 |
| | 5XUS[16] (LbCas12a, 20 bp) | 7.7 |
| S4b | 6GTD[1] (FnoCas12a, I2) | 9.7 |
| | 6GTE[1] (FnoCas12a, I3) | 4.3 |
| | 6GTG[1] (FnoCas12a, I4) | 6.5 |
| | 5NFV[9] (FnoCas12a, 20 bp) | 5.7 |

| | | |
|---|---|---|
| | 6I1L [15](FnoCas12a, 20 bp) | 7.5 |
| | 6I1K[15] (FnoCas12a, 20 bp) | 6.5 |
| | 8SFL[10] (AsCas12a, 15 bp) | 6.5 |
| | 8SFN[10] (AsCas12a, 16 bp) | 8.7 |
| | 8SFO[10] (AsCas12a, 20 bp) | 8.3 |
| | 8I54[12] (LbCas12a, 14 bp) | 13.3 |
| | 5XUS[16] (LbCas12a, 20 bp) | 6.9 |

**Supplementary Table 3: Movement of REC2 and Nuc domains.** The distance between residues D468 (REC2) and V1102 (Nuc) highlights the movement between the REC2 and Nuc domains, illustrating the closed-to-open conformational transition of the Cas12a complex.

| **State** | **Distance (Å)** |
|---|---|
| S1 | 10 |
| S2 | 14 |
| S3 | REC2 missing |
| S4a | 46 |
| S4b | 31 |
| 6GTC (I1) | 8 |
| 6GTE (I3) | 38 |
| 6GTG (I4) | 340 |

**Supplementary Table 4**: **Distance between the REC1 and REC2 domains between the different states.** The distance between the residues K85 (REC1, HLH) and C473 (REC2) were measured to analyze the movement of REC1 and REC2 domains.

| **State** | **Distance (Å)** |
|---|---|
| S1 | 40 |
| S2 | 53 |
| S3 | REC 2 missing |
| S4a | 77 |
| S4b | 71 |
| 6GTC (I1) | 35 |
| 6GTD (I3) | 70 |
| 6GTG (I4) | 63 |
| S1 and S2 | 13 |
| S1 and S4a | 37 |
| S1 and S4b | 31 |
| S2 and S4a | 24 |
| S2 and S4b | 18 |

**Supplementary Table 5: (A) Movement of the REC1 domain with respect to the NUC lobe.** Translation and rotation angle between different states were measured using a triangle passing through the Cα-atom of L621 in the PI domain, F1124 in the Nuc domain, and N93 (residue of HLH segment) in the REC1 domain.

| State | PI to Nuc (Å) | PI to REC1 (Å) | Nuc to REC1 (Å) | Angle between REC1 and NUC-lobe axis (°) |
|---|---|---|---|---|
| S1 | 63 | 57 | 85 | 43 |
| S2 | 58 | 71 | 83 | 57 |
| S3 | 58 | 70 | 85 | 55 |
| S4a | 58 | 72 | 86 | 56 |
| S4b | 58 | 69 | 85 | 53 |
| I1 | 64 | 55 | 81 | 43 |
| I3 | 60 | 67 | 81 | 54 |
| I4 | 58 | 58 | 81 | 46 |
| Difference between the states | | | | |
| S1 to S2 | -5 | 14 | -2 | 14 |
| S2 to S3 | 0 | -1 | 2 | -2 |
| S3 to S4a | 0 | 2 | 1 | 1 |
| S3 to S4b | 0 | -1 | 0 | -2 |
| S1 to S3 | -5 | 13 | 0 | 12 |
| S1 to S4a | -5 | 15 | 1 | 13 |
| S1 to S4b | -5 | 12 | 0 | 10 |
| I1 to I3 | -4 | 12 | 0 | 11 |
| I1 to I4 | -6 | 3 | 0 | 3 |
| S4a to I4 | 0 | -14 | -5 | -10 |
| S4b to I4 | 0 | -11 | -4 | -7 |
| I3 to I4 | -2 | -9 | 0 | -8 |

**Supplementary Table 5: (B) Movement of the REC2 domain with respect to the NUC lobe.** Translation and rotation angle between different states were measured using a triangle passing through Cα-atom of L621 in the PI domain, F1124 in the Nuc domain, and K539 of REC2 domain. The distance was not measured for S3 as it lacked the REC2 domain.

| State | PI to Nuc (Å) | PI to REC2 (Å) | Nuc to REC2 (Å) | Angle between REC2 and NUC-lobe axis (°) |
|---|---|---|---|---|
| S1 | 85 | 57 | 50 | 35 |
| S2 | 83 | 64 | 49 | 36 |
| S4a | 86 | 69 | 63 | 53 |
| S4b | 85 | 63 | 54 | 47 |
| I1 | 81 | 60 | 46 | 47 |
| I3 | 81 | 66 | 55 | 53 |
| I4 | 81 | 72 | 55 | 60 |
| | Difference between the states | | | |
| S1 to S2 | -1 | 7 | -1 | 1 |
| S2 to S4a | 2 | 5 | 14 | 17 |
| S2 to S4b | 2 | -1 | 5 | 11 |
| S1to S4a | 1 | 12 | 13 | 18 |
| S1 to S4b | 0 | 6 | 4 | 12 |
| I1 to I3 | 0 | 6 | 9 | 6 |
| I1 to I4 | 0 | 12 | 9 | 13 |
| S4a to I4 | -5 | 3 | -8 | 7 |
| S4b to I4 | -4 | 9 | 1 | 13 |
| I3 to I4 | 1 | 6 | 0 | 7 |

**Supplementary Table 6: Nucleic acid base pairing and visibility in different states**

| State | Number of nucleotides visible | | | Number of nucleotides base pairing |
|---|---|---|---|---|
| | crRNA | TS DNA | NTS DNA | crRNA:TS DNA |
| S1 | 27 | - | - | - |
| S2 | 28 | 8 | - | 8 |
| S3 | 30 | 11 | 7 | 8 |
| S4a | 36 | 16 | 8 | 11 |
| S4b | 31 | 16 | 9 | 9 |

**Supplementary Table 7: Bending of BH.** The angle between BH and nucleic acid in different states were measured using a triangle passing through the Cα-atom of D/P 970 in the loop region, Y953 in the beginning of BH, and C4′ of dT-8 on the TS DNA. (Note: due to lack of DNA in the binary structure we used DNA from 6GTC (RMSD: 1.15 Å) to measure the distance of BH form the nucleic acid).

| **States** | D/P970-Y953 (Å) | Y953- dT-8 (Å) | D/P970-dT-8 (Å) | Angle of BH shift (°) |
|---|---|---|---|---|
| 6GTC | 21 | 24 | 16 | 80 |
| S1 | 22 | 25 | 16 | 81 |
| S2 | 20 | 24 | 15 | 85 |
| S4a | 24 | 26 | 21 | 70 |
| S4b | 26 | 21 | 20 | 52 |
| 6GTE | 28 | 23 | 24 | 52 |
| 6GTG | 24 | 19 | 15 | 52 |
| 6GTC and 6GTG | 3 | -5 | -1 | -28 |
| S1 and S4a | 2 | 1 | 5 | -11 |
| S1 and S4b | 4 | -4 | 4 | -29 |
| S2 and S4a | 4 | 2 | 6 | -15 |
| S2 and S4b | 2 | -5 | 0 | -33 |
| S4a and 6GTG | 0 | -7 | -6 | -18 |
| S4b and 6GTG | -2 | -2 | -5 | 0 |
| 6GTE and 6GTG | -4 | -4 | -9 | 0 |

**Supplementary Table 8**: **Interaction of BH with RNA:DNA hybrid.** We measured the distance between the BH (K/P 969 Cα-atom) and RNA:TS DNA hybrid (C4'- atom). The position of nucleic acid is mentioned in parentheses.

| **States** | **BH -R-loop (Å)** |
|---|---|
| 5NG6 (Binary) | 34 (U4) |
| S1 | 34 (A4) |
| S1 | 21 (C7) |
| S2 | 18 (dT-8) |
| 6GTC | 17 (dT-8) |
| 6GTD | 17 (dT-8) |
| 6GTE | 23 (dT-8) |
| 6GTG | 14 (dT-8) |
| S4a | 21 (dT-8) |
| S4b | 20 (dT-8) |
| 6GTG | 6 (U11) |
| S4a | 20 (U11) |
| S4b | 27 (U11) |

**Supplementary Table 9**: **Positional shift of HLH domain.** The distance between Cα-atom of N93 of different states measured to analyze the positional shift of the HLH domain across the different states.

| **States** | **Distance of N93 between two states (Å)** |
|---|---|
| S1 and S2 | 17 |
| S2 and S3 | 8 |
| S3 and S4a | 4 |
| S3 and S4b | 0.7 |
| S2 and S4a | 5 |
| S2 and S4b | 7 |
| S1 and S4b | 14 |
| S1 and I1 | 4 |
| S1 and I4 | 12 |
| S2 and I4 | 16 |
| S3 and I2 | 5 |
| S3 and I3 | 3 |
| S3 and I4 | 7 |
| S4a and I4 | 11 |
| S4b and I3 | 3 |
| S4b and I4 | 8 |
| I1 and I4 | 12 |
| I2 and I4 | 13 |
| I3 and I4 | 6 |
| I3 and S4b | 2 |

**Supplementary Table 10: Movement of REC-linker and finger helix.** Distance between REC-linker (K326) and crRNA (U6) and finger helix (N301) and TS (dT-14) DNA. Distance is absent for the states lacking density for dT-14.

| State | K326:U6 (Å) | N301:dT-14 (Å) |
|---|---|---|
| S1 | 19.4 | - |
| S2 | 6.3 | - |
| S3 | 3.1 | - |
| S4a | Linker density missing | 15.2 |
| S4b | 2.6 | 20.8 |
| 6GTC (I1) | 15.5 | - |
| 6GTD (I2) | 12.2 | - |
| 5NFV | 6.4 | 9.0 |
| 6GTG (I4) | 2.7 | 4.1 |
| 5NG6 (binary) | 18.4 | - |

**Supplementary Table 11: Oligonucleotide sequences used for FnoCas12a$^{KD2P}$ complexing.** The sequence of crRNA constructs used for *in vitro* transcription consists of the T7 promoter sequence used to facilitate *in vitro* transcription. T7 promoter is in italics, bold-upper case sequence represents guide region and the nucleotides underlined represents PAM sequence.

| **Oligo name** | **Sequence** |
|---|---|
| MCH TS (24-nt) | 5'- **GCCTTATTAAATGACTTCTC**TAAA -3' |
| MCH NTS (24-nt) | 5'- TTTA**GAGAAGTCATTTAATAAGGC** -3' |
| crRNA template strand | 5'- AGTG**GCCTTATTAAATGACTTCTC**ATCTACAACAGTAGAAATT*CCCTATAGTGAGTCGTATTAATTTC* -3' |
| T7 promoter strand | 5'- *GAAATTAATACCACTCACTATAGGG* -3' |
| crRNA | 5'- AAUUUCUACUGUUGUAGAU**GAGAAGUCAUUUAAUAAGGC**CACU -3' |
